# *In Vivo* Study of Key Transcription Factors in Muscle Satellite Cells by CRISPR/Cas9/AAV9-sgRNA Mediated Genome Editing

**DOI:** 10.1101/797746

**Authors:** Liangqiang He, Yingzhe Ding, Yu Zhao, Karl K. So, Xianlu L. Peng, Yuying Li, Jie Yuan, Hao Sun, Huating Wang

## Abstract

Skeletal muscle satellite cells (SCs) are adult muscle stem cells responsible for injury induced muscle regeneration. Despite advances in the knowledge of molecular mechanisms regulating SC lineage progression, our understanding of key transcription factors (TFs) and their regulatory functions in SCs in particularly the quiescent and early activation stages remains incomplete due to the lack of efficient method to screen and investigate the stage-specific key TFs. In this study, we succeeded in defining a distinct list of key TFs in early stages of SC fate transition using the paradigm of super enhancers (SEs). Particularly, leveraging the Cre-dependent Cas9 knockin mice and AAV9 mediated sgRNAs delivery, we generated a facile muscle specific genome editing system which allows gene depletion in SCs *in vivo.* Using *MyoD* locus as a proof of concept, we demonstrated that this CRISPR/Cas9/AAV9-sgRNA system can efficiently introduce mutagenesis at target locus and recapture the phenotypes reported in knockout mice. Further application of the system on key TFs, *Myc, Bcl6* and *Pknox2,* revealed their distinct functions in the early stage of SC activation and damage induced muscle regeneration. Altogether our findings have proven the CRISPR/Cas9/AAV9-sgRNA system as a robust way for *in vivo* genome editing and elucidation of key factors governing SC activities.

## INTRODUCTION

Skeletal muscle possesses excellent regeneration potential after injury, which is executed mainly by adult muscle stem cells also called satellite cells (SCs). In healthy adult skeletal muscle, SCs surrounded by myofiber sarcolemma and basal lamina(1) are maintained in a quiescent state, characterized by paired-box gene 7 (Pax7) expression. Upon traumas, these dormant SCs quickly activate and re-enter the cell cycle to give rise to myoblasts characterized by elevation of two basic helix-loop-helix (bHLH) transcription factors myogenic factor 5 (Myf5) and MyoD. Then a large portion of myoblasts down-regulate Pax7 expression and undergo myogenic differentiation to form new muscle fibers by inducing Myogenin and myogenic regulatory factor 4 (MRF4) expression. Meanwhile, a sub-population of activated SCs retaining high Pax7 expression return to quiescence to restore the SC pool(1,2).

Transcriptional regulations orchestrated by transcription factors (TFs) represent the key mechanisms driving SC fate transition and lineage progression. A wealth of knowledge has been accumulated in our understanding of TF regulation during myoblast differentiation into myotube. In addition to myogenic specific TFs such as the above mentioned MyoD, Myogenin and MRF4, ubiquitously expressed TFs such as YYi also execute important regulatory roles in controlling myogenic differentiation(3–5). In comparison, our knowledge in transcriptional regulation driving the early stages of SC activity remains limited. As SC fate determinant, Pax7 is critical to maintain SC quiescence and its deficiency results in progressive loss of the SC pool mainly owing to failed proliferation and precocious differentiation(6,7). It has also been shown that knockout of Pax7 in muscle progenitor cells induces a cell fate switch to brown adipocytes(8). On the other hand, MyoD is known as the marker of activated SCs and crucial for myogenic lineage progression(1,9–11). In addition to Pax7 and MyoD, TFs from Notch pathway including Rbpj and those in the Hes and Hey families also play crucial roles in coordinating SC quiescence and activation(1). Knockout of *Rbpj* or double deletion of *Hey1* and *HeyL* in SCs induces spontaneous activation and depletion of SC pool(12,13). Stra13, a bHLH family TF, has also been reported to coordinate SC activation by antagonizing Notch signaling(14). Despite isolated studies on TF regulation of SC quiescence and early activation, a systematic approach allowing identification of potentially functional key TFs is needed. We reason this goal can be achieved based on recent advances in our understanding of enhancer regulation. Enhancer is a DNA element that can activate gene transcription in a distance- and orientation-independent manner(15). A subset of enhancers termed super enhancers (SEs) have been defined as large DNA regions with unusual enriched binding of key TFs, co-activators, and Mediator (Med1)(16). According to our recent findings, during myoblast differentiation, SE activation is orchestrated by key TFs such as MyoD, Myogenin, FoxO3 and these TFs are themselves regulated by SEs(17). Such inter-connected regulatory circuits can thus be exploited to identify key TFs. For example, Saint-André *et al* succeeded in generating cell-type specific core transcriptional regulatory circuitries (CRCs) for 75 human cell and tissue types by leveraging this unique feature of SEs(18).

The downstream functional dissection of identified TFs can be greatly facilitated by the quickly matured CRISPR/Cas9 mediated loss-of-function tools. By directing of a single guide RNA (sgRNA), Cas9 is able to modify genome at a specific site(19). The ease of CRISPR/Cas9 allows genome editing in many cell lines and various species. Recently, its *in vivo* application in mouse is emerging to generate mouse models and correct genetic diseases by viral or non-viral based Cas9/sgRNA delivery(20). In particular, several reports described its utility in skeletal muscle tissue, treating Duchenne muscular dystrophy (DMD), a X-chromosome-linked neuromuscular disorder caused by mutation of the dystrophin gene(21). In these attempts, adeno-associated virus (AAV) mediated Cas9/sgRNA delivery was employed to delete the mutated exon so that forming a truncated dystrophin protein with partial function in the *mdx* mouse model of DMD(22–24). Normally co-transduction of multiple AAV vectors is required to deliver the Cas9 and sgRNAs separately considering the restricted capacity of AAV virus (~4.7 kb)(22–24), which limited the modification efficiency since successful editing only occurs in nuclei simultaneously receiving all the components. In addition, almost all of the genomic modifications were done in post-mitotic myofibers and it remains to be determined whether such AAV/Cas9/sgRNA based tool can be applied to edit SCs *in vivo*(21), which is more attractive since it provides a sustained reservoir of cells to generate dystrophin expressing myofibers during normal muscle turnover(21). While it is known that SCs are resistant to AAV transduction(21,25,26), one study reported successful editing in SCs with modest efficiency by injecting the AAV virus at neonatal stage(24) and the same group recently reported efficient transduction of multiple AAV serotypes to SCs in adult mice by a Cre/lox fluorescent reporter tracking system(27). Nevertheless, whether quiescent SCs (QSCs) in adult stage are permissive to CRISPR/Cas9 mediated genome editing is still unknown.

Here, in this study, we first identified lists of key TFs in quiescent, early activating SCs and activated myoblasts using the paradigm of SEs. The unsuccessful attempts of exploring their functions in isolated SCs led us to resort to *in vivo* genome editing tools. We thus generated a muscle restricted genome editing system which allows efficient gene depletion in SCs by integrating the Cre-dependent Cas9 knockin mice with AAV9 mediated sgRNA delivery in postnatal stage. First, we proved that expression of Cas9 in SCs and skeletal muscle had no obvious impact on SC homeostasis and muscle regeneration. Next, using *MyoD* locus as a paradigm, we demonstrated the high efficiency of this system in deleting MyoD in SCs, which expectedly impaired myoblast differentiation. Further applying this system to edit key TFs, *Myc, Bcl6* and *Pknox2,* led to successful decrease of the TFs in SCs which caused distinct effects on SC activation and muscle regeneration. Additionally, unsuccessful attempts to edit QSCs in adult mouse led us to conclude that QSCs are refractory to CRISPR/Cas9 mediated genome modification even with efficient AAV9 transduction. Altogether our studies have established a CRISPR/Cas9/AAV9-sgRNA mediated system for *in vivo* genome editing and elucidation of key TF functions in regulating SC activities.

## MATERIALS AND METHODS

### Mice

All animal experiments were performed according to guidelines for experimentation with laboratory animals set in institutions and approved by the CUHK Animal Ethics Committee. Pax7-nGFP, Pax7_Cre_ and Pax7_CreER_ mice were kindly provided by Dr. Zhenguo Wu (Hong Kong University of Science & Technology). The Cre dependent Rosa26_Cas9-EGFP_ knockin mice (B6;129-Gt(ROSA)26Sor_tm1(CAG-cas9*-EGFP)Fezh/J_; stock number 024857) were obtained from the Jackson Laboratory. To generate skeletal muscle specific Cas9 knockin mice, homozygous Pax7_Cre_ or Pax7_CreER_ mice were crossed with Rosa26_Cas9-EGFP_ mice. For inducible Cas9 expression, Tamoxifen (100 mg/Kg body weight) was injected intraperitoneally into Pax7_ER-Cas9_ mice for five consecutive days. Each mouse strain was genotyped by PCR using DNA extracted from mouse tail tissues. To induce acute injury, ~8-week-old mice were injected with 50 μL of 1.2% BaCl_2_ (w/v in H_2_O) solution into the tibialis anterior (TA) muscle. TA muscles were harvested at designated time points for further analysis. Primers used for genotyping are listed in Supplementary Data S1.

### Cell culture

Mouse C2C12 myoblast cells (CRL-1772) and human HEK293FT cells were obtained from ATCC and cultured in DMEM with 10% FBS, 100 units/ml penicillin, 100 μg/ml streptomycin and 2 mM L-glutamine (GM, growth medium) in 5% CO_2_ at 37 °C.

### Satellite cell isolation, culture, transfection and EdU incorporation assay

Satellite cells were isolated from Pax7-nGFP, Pax7_Cas9_ or Pax7_ER-Cas9_ mice by FACS based method on GFP signal as reported previously(28). Isolated satellite cells were cultured in growth medium including F10 medium (Merck Millipore) with 20% FBS, 1 % penicillin/streptomycin and 5 ng/mL basic fibroblast growth factor (bFGF) at 37°C in 5% CO_2_. To induce spontaneous differentiation, satellite cells were cultured in growth medium for two or four days. For *in vitro* transfection, satellite cells isolated from Pax7-nGFP mice were transfected with siRNAs or plasmids two hrs after seeding using Lipofectamine 3000 reagent according to the manufacturer’s instructions. SiRNA oligos against selected TFs were from GenePharma (Shanghai, China). For transient transfection, the final concentration of siRNAs was 100 nM. For EdU incorporation assay, EdU was added to the SC culture medium with a final concentration of 10 mM and the incorporation assay was detected by Click-iT EdU kit (Invitrogen) according to the manufacturer’s instructions. Related sequences for each siRNA are listed in Supplementary Data S1.

### Plasmids

To generate overexpression plasmids for the selected TFs, the cDNA of each TF was amplified by PCR and inserted into the pRK5-flag vector (kindly gift from Dr. Jing Zhang(29)) using BamH I and Hind III sites (Xho I and Hind III sites for Pknox2). To construct the AAV9-sgRNA transfer plasmids, the AAV: ITR-U6-sgRNA (backbone)-pCBh-Cre-WPRE-hGHpA-ITR (Addgene, 60229) was used as donor plasmid. Coding sequencing for DsRed was PCR-amplified and cloned into the donor plasmid through replacing the sequence encoding Cre using Age I and EcoR I sites. The pCBh promoter was substituted with CMV promoter to drive DsRed expression. For single sgRNA expression system, sgRNA with the highest editing efficiency by Surveyor nuclease assay was inserted into the AAV9-sgRNA vector (AAV: ITR-U6-sgRNA(backbone)-CMV-DsRed-WPRE-hGHpA-ITR) using Sap I site. To generate dual AAV-sgRNAs expression plasmid, the second sgRNA with good editing efficiency and targeting about 100 bp-300 bp away from the first sgRNA was constructed into the AAV9-sgRNA vector together with the gRNA cassette and U6 promoter using Xba I and Kpn I sites. To increase the editing efficiency, the two selected sgRNAs are designed to target 5’ end of the coding region and the predicted deletion should cause frameshift of the gene. Primers for AAV9-sgRNA vector construction are listed in Supplementary Data S1.

### SgRNA design, selection and Surveyor nuclease assay

Surveyor nuclease assay was performed according to the published protocol(30). In brief, site specific sgRNAs were predicted following a web tool Crispor(31) (http://crispor.tefor.net/). To minimize off-target effects, only sgRNAs with a score higher than 5 were selected. To determine the editing efficiency, annealed oligonucleotides were constructed into a Cas9-EGFP expressing vector (pX458, Addgene) using Bbs I site. SgRNA-pX458 plasmids were transiently transfected into C2C12 using Lipofectamine 3000. GFP positive cells were sorted out using FACS 48 hrs after transfection and cultured for another two days. Genomic DNAs were extracted by QuickExtract (Epicentre) solution and amplified by Phusion High-Fidelity DNA Polymerase (NEB) using primers against the edited locus. Purified PCR products were subject to Surveyor nuclease assay (Integrated DNA Technologies) according to the manufacturer’s protocol. Briefly, 360 ng of DNA was denatured at 95°C and re-annealed to form hetero-duplexes. The resulting product was digested by Surveyor Nuclease S at 42 °C for 30 minutes and separated on a 2% agarose gel. The percentage of indel formation was determined by relative band intensities. Sequences for sgRNAs and primers used for Surveyor assay are listed in Supplementary Data S1.

### AAV9 virus production, purification and injection

AAV9 virus particles were produced in HEK293FT cells by the triple transfection method(32). In brief, HEK293FT cells were seeded in T75 flask and transiently transfected with AAV9-sgRNA vector (5 μg), AAV9 serotype plasmid (5 μg), and pDF6 (AAV helper plasmid) (10 μg) at a ratio of 1:1:2 using polyethyleneimine (PEI) when the cell reached 80%~90% confluent. Twenty-four hrs after transfection, the cells were changed to growth medium (DMEM with 10% FBS, 100 units/ml penicillin, 100 μg/ml streptomycin and 2 mM L-glutamine) and cultured for another forty-eight hrs. The cells were harvested by Trypsin-EDTA (Gibco) and washed with PBS for two times. To release the AAV9 virus, the pellet was re-suspended with lysis buffer (Tris-HCl, PH 8.0, 50 mM; NaCl, 150 mM) followed by three sequential freeze-thaw cycles (liquid nitrogen/37°C). The lysate was treated with Benzonase (Sigma) together with MgCl_2_ (final concentration: 1.6 mM) at 37°C for 0.5~1 hr followed by centrifugation at 3,000 rpm for 10 minutes. The supernatant was filtered with 0.45 μm sterile filter and added with equal volume of 1 M NaCl and 20% PEG8000 (w/v) to precipitate the virus at 4 °C overnight. After spinning the mixture at 12,000 g, 4 °C, for 30 minutes, the supernatant was discarded and the pellet was re-suspended with sterile PBS and then subject to centrifugation at 3,000 g for 10 minutes. Equal volume of chloroform was then added and shaken. The mixture was spun down at 12,000 g for 15 minutes at 4 °C. The aqueous layer was filtered by 0.22 μm sterile filter and passed through a 100 kDa MWCO (Millipore). The concentrated solution was washed with sterile PBS for three times. The tilter of the AAV9 virus was determined by qRT-PCR using primers targeting the CMV promoter. AAV9 serotype plasmid and pDF6 are kind gifts from Dr. Bin Zhou(33). For AAV9 administration, 50-100 μL of AAV9-sgRNA (high dose: 5×10_11_ vg; middle dose: 1 × 10_11_ vg; low dose: 0.2×10_11_ vg) or control (AAV9-sgRNA vector without sgRNAs insertion) virus was diluted in saline and injected systemically through intraperitoneal (IP) injection on postnatal day 2 (P2) or locally to the skeletal muscles through intramuscular (IM) injection on P10. All the used primers are listed in Supplementary Data S1.

### Immunoblotting, immunostaining and immunohistochemistry

Tissue samples were homogenized in 400 μl of ice cold RIPA buffer with a protease inhibitors cocktail (Sigma-Aldrich) and lysed on ice for 40 minutes. Total cell extracts for Western blot were prepared as described previously(3,5). Following antibodies were used: MyoD (Dako, M3512, 1:2000), α-Tubulin (Santa Cruz Biotechnology, sc-23948, 1:2000), Cas9 (Cell Signaling Technology, #14697, 1:1000), Pax7 (Developmental Studies Hybridoma Bank; 1:1000), GFP (Santa Cruz Biotechnology, sc-8334, 1:1000), GAPDH (Santa Cruz Biotechnology, sc-137179, 1:2000), Esr1(Santa Cruz Biotechnology, sc-8005, 1:1000), Myc (Santa Cruz Biotechnology, sc-40, 1:1000), Bcl6 (Santa Cruz Biotechnology, sc-365618, 1:500), Pknox2 (Santa Cruz Biotechnology, sc-101857, 1:1000), Sox4 (Santa Cruz Biotechnology, sc-518016, 1:1000), Rora (Invitrogen, PA5-23268, 1:1000), Myogenin (Santa Cruz Biotechnology, sc-576, 1:1000). For immunofluorescence staining, following antibodies and related dilutions were used: MyoD (Dako, M3512, 1:800), Pax7 (Developmental Studies Hybridoma Bank; 1:100), DsRed (Santa Cruz Biotechnology, sc-390909, 1:400). Hematoxylin and eosin (H&E) staining on frozen muscle sections was performed as previously described(34). Immunofluorescence staining on frozen muscle sections was performed using the following antibodies: Laminin (Sigma, L9393, 1:800), eMyHC (Sigma, 1:200), Pax7 (Developmental Studies Hybridoma Bank; 1:50), MyoD (Dako, M3512, 1:200). All fluorescent images were captured with a fluorescence microscope (Leica).

### qRT-PCR

Total RNAs from cells were extracted using TRIzol reagent (Life Technologies) according to the manufacturer’s instructions and cDNAs were prepared using PrimeScriptTM RT Master Mix kit (Takara, RR036A). Analysis of mRNA expression was performed with SYBR Green Master Mix (Life Technologies) on a 7900HT System (Life Technologies). All the used primers are listed in Supplementary Data S1.

### RNA-seq and data analysis

RNA-seq was performed as described previously(5,35). Total RNAs were extracted from freshly isolated or cultured SCs and subject to poly(A) selection (Ambion, 61006) followed by library preparation using NEBNext® Ultra™ II RNA Library Preparation Kit (NEB). Libraries with barcodes were pooled at equal concentrations and sequenced on the Illumina HiSeq 1500 platform. For data analysis, Sequenced reads were mapped to reference mouse genome using TopHat (v2.0.13). Cufflinks (v2.1.1) was then employed to estimate transcript abundance. Abundance was reported in Fragments Per Kilobase per Million (FPKM). Differentially expressed genes were identified if the change of expression level exceeds a fold change threshold (> 2).

### ChIP-seq and data analysis

ChIP-seq and the data analysis were performed following the procedures described in our previous studies(3,36,37). In brief, chromatins from freshly isolated or cultured SCs were fragmented using sonicator and incubated with 5 μg antibody against histone H3-K27 acetylation (Abcam, ab4729, rabbit poly-clonal) overnight at 4 °C. The purified DNA (200 ng) was subject to library preparation using NEBNext® Ultra™ II DNA Library Preparation Kit (NEB). Libraries with barcodes were pooled and sequenced on the Illumina HiSeq 1500 platform. For data analysis, a standard approach was used to conduct base calling and convert the results into raw reads in FASTQ format. After adapter trimming and quality filtering, reads were first aligned to mm9 reference genome by Bowtie2 with default parameters, then removed duplication and called enriched regions (peaks) by MACS2 with q-value equal to 0.01. Defining of enhancers and super enhancers was performed as described before with minor adjustment(17). Enriched H3K27ac regions(peaks) were identified by MACS2 using merged bam files from all replicates at each stage with q-value 0.01. Peaks were subject to a filter to exclude the ENCODE blacklisted regions as well as those within +/-2 kb of a Refseq Transcription Start Site (TSS). The filtered peaks were defined as enhancers and assigned to expressed transcripts (RPKM >0.5) whose TSSs are nearest to the center of the enhancer regions. The enhancers within 12.5 kb of one another and can be assigned to the same genes were stitched together and subject to the ROSE algorithm for super enhancer identification(38). SE constituents were extended by 500 bp on both side and used as input regions to scan motifs for transcription factors by FIMO with default parameters. The Position weight matrix (PWM) of the motifs were obtained from the TRANSFAC database.

### Deep-seq

Deep-seq was performed as previously described(39–41). Briefly, genomic DNAs from AAV9-sgRNA infected SCs were amplified using Q5 High-Fidelity 2× Master Mix (NEB). PCR products were purified through QIAQuick PCR purification kit (Qiagen) and subject to library preparation using NEBNext® Ultra™ II DNA Library Preparation Kit (NEB). Libraries with barcodes were pooled and sequenced on the Illumina HiSeq 1500 platform. For data analysis, an online tool CRISPResso2 (http://crispresso.pinellolab.partners.org/)(42) was employed to calculate the indel occurrence. Primers used for Deep-seq are listed in Supplementary Data S1.

### Statistical analysis

All the data are presented as the mean + standard error of the mean (sd). Statistical comparison of two groups was conducted using Student’s t-test and the p values presented in the Figures are \**P* < 0.05, \*\**P* < 0.01 and \*\*\**P* < 0.001.

## RESULTS

### Prediction of key transcription factors regulating SC quiescence and activation

Despite advances in the knowledge of molecular mechanisms regulating SC lineage progression, our understanding of key TFs and their functions in SCs in particularly the quiescent and early activating stages remains incomplete. In this study, we thus set out to explore functionality of key TFs in early stages of SC fate transition. We started by defining the TFs that are key to cell identity of QSCs and activated SCs (ASCs or myoblasts) by leveraging the unique trait of SEs. To this end, we performed H3K27ac ChIP-seq in freshly isolated SCs (FISCs) from Pax7-nGFP mice(43) and ASCs cultured in growth medium for 24 hrs (Figure 1A, Supplementary Figure S1A and B). Identification of enhancer constituents was conducted following the standard analysis pipeline(17,37) using intergenic and intronic H3K27ac peak regions (Supplementary Figure S1C and Supplementary Data S2). A high portion of enhancers were exclusively present in FISC or ASC stage (Supplementary Figure S1D and Supplementary Data S3), demonstrating the stage specificity and the remodeling of enhancer landscape during SC activation. A slightly modified ROSE algorithm(16,17) was then applied to define SEs in both FISCs and ASCs (Supplementary Figure S1E and F) leading to the identification of a total of 57 and 163 SEs (Supplementary Data S4), respectively. Based on the prior knowledge that key TFs are generally associated with SEs and tend to occupy their own and other key TFs’ SEs to form interconnected auto-regulatory loops(16,18,44), we adopted a pipeline(18) to predict key TFs in FISCs and ASCs (Figure 1B). In FISCs, a total of 650 active TF genes were identified among which 220 are enhancer-associated while 15 are SE-associated; 12 auto-regulated TFs were further identified and 10 were determined as key TFs based on the activity and expression level of each candidate (Supplementary Data S5); the list includes Pax7, Runx1, Mef2d, Cebpb, Klf9, Sox4, Esr1, Six1, Bcl6 and Rora (Figure 1B and Supplementary Figure S1E). Similarly, in ASCs, a total of 11 key TFs (Supplementary Data S5) including Pax7, Tgif1, Runx1, Nfix, MyoD, Six1, Cebpb, Myc, Nfia, Pknox2 and Bcl6 were defined (Figure 1B and Supplementary Figure S1E). Consistent to previous studies, these key TFs indeed displayed higher expressions compared to other SE or enhancer associated genes (Supplementary Figure 1G and Supplementary Data S6). Of note, Pax7 is presented in both QSC and ASC lists while MyoD only appears in ASC list (Supplementary Figure S1E and H), which is consistent with the well-known functions of these two TFs in orchestrating SC homeostasis and activation. On the other hand, Cebpb, Bcl6, Six1, Runx1 were identified in both FISCs and ASCs. Considering the functions of these TFs in regulating SC fate transition remain largely unknown, we reasoned the downstream investigation of their specific roles *ex vivo* and *in vivo* will lead to unprecedented discovery of novel aspects of transcriptional regulation in SCs.

**Figure 1.**
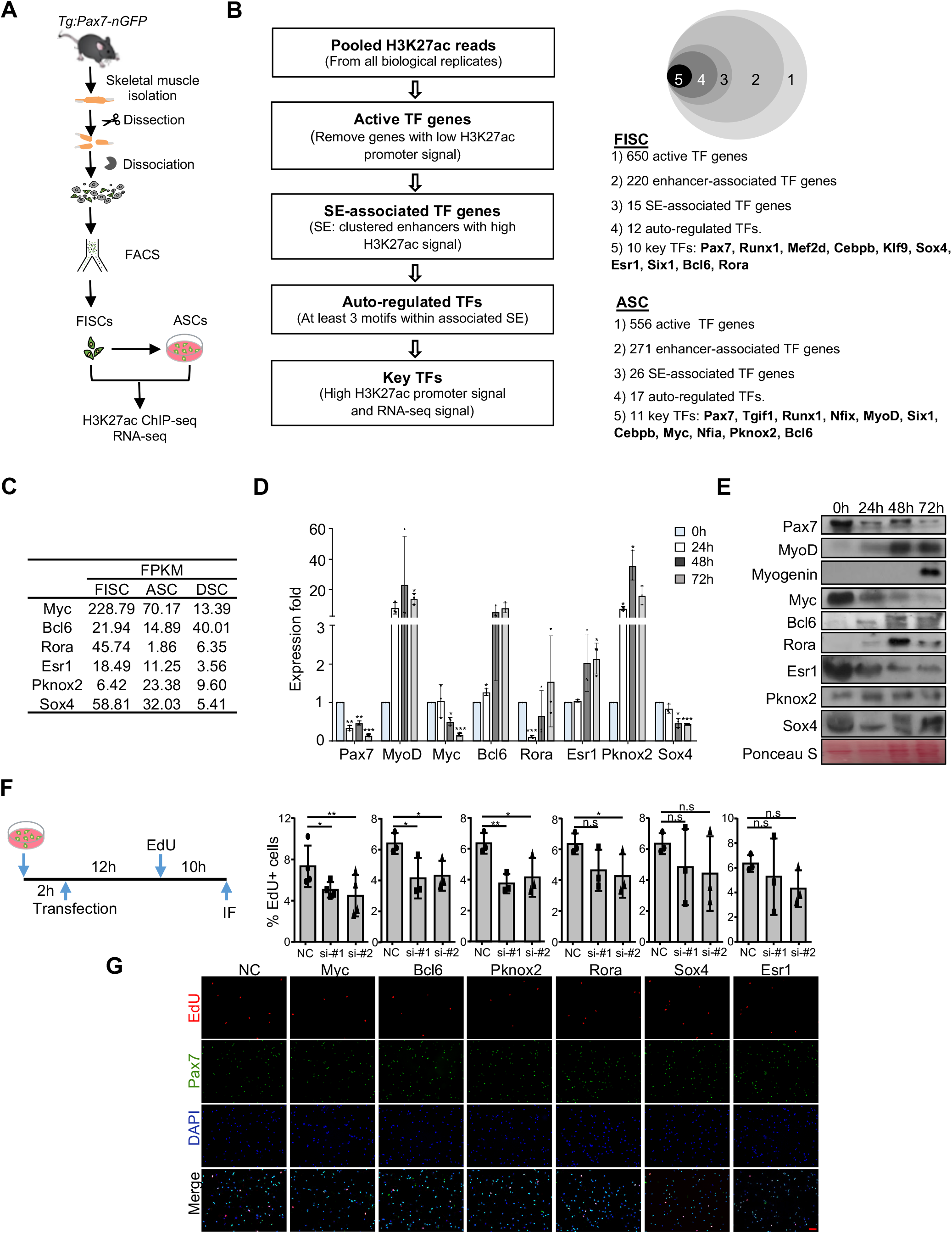
Prediction and *ex vivo* testing of key transcription factors regulating SC quiescence and activation. (**A**) Schematic illustration of the protocol for SC isolation from Pax7-nGFP mice. (**B**) Schematic illustration of the bioinformatics pipeline for predicting key TFs in FISC and ASC. (**C**) Expression of selected key TFs in FISC, ASC and DSC examined by RNA-seq. FPKM, fragments per kb of transcript per million mapped reads. (**D**) Dynamic expression of selected TFs in SCs cultured for 0, 24, 48 and 72 hrs determined by qRT-PCR (n = 3 in each group). (**E**) The protein levels of the above selected TFs were determined by Western blot. Ponceau S staining of the member was used as loading control. (**F**) Left: schematic illustration of the EdU incorporation assay to examine the change of SC activation after knockdown of each key TF. Right: the percentage of EdU positive cells was quantified from at least three experiments and presented as mean ± s.d. (**G**) Representative images to show EdU, Pax7 staining of the above cells. Scale bar, 75 μm. All qRT-PCR data were normalized to 18S mRNA and presented as mean ± s.d. *P<0.05, **P<0.01, ***P<0.001. ns, no significance.

### *Ex vivo* testing of selected key TFs

Based on their expression levels and known relevancy in muscle cells, six key TF candidates, including Myc, Bcl6, Rora, Esr1, Pknox2, Sox4 were selected for further functional investigation *ex vivo* (Figure 1C). Briefly, Myc has been reported to inhibit C2C12 differentiation through preventing myoblast fusion(45) whereas Bcl6 plays a pro-differentiating role by restraining apoptotic cell death(46). Sox4 is a member of the Sox TFs family that facilitates differentiation of skeletal myoblasts through regulating caldesmon expression(47). Esr1, a ligand-activated TF, also known as estrogen receptor-α, is required for estradiol-stimulated proliferation of cultured bovine SCs(48). Although nothing is known on Pknox2 function in SCs, one of its paralog, Pknox1, can regulate mitochondrial oxidative phosphorylation in skeletal muscle(49). Rora, a member of nuclear hormone receptors family, accelerates myoblast differentiation by directly interacting with MyoD(50).

Based on RNA-seq measurement, the expressions of these six TFs displayed varied dynamics during SC lineage progression (Figure 1C). *Myc* exhibited the highest expression in FISCs, which continued to decrease in ASCs and differentiated SCs (DSCs); mRNA expressions of *Esr1* and *Sox4* also displayed continuous decrease. *Bcl6* expression decreased in ASCs vs. FISCs but increased dramatically in DSCs, whereas *Rora* presented a sharp decrease in ASCs vs. FISCs, which bounced back slightly in DSCs. *Pknox2,* on the other hand, showed increased expression from FISCs to ASCs, which then decreased in DSCs. To validate the results, by qRT-PCR, we further examined their expression in SCs cultured for 24, 48 and 72 hrs, during which period SCs activated (24 hr), proliferated (48 hr) and differentiated (72 hr) (Figure 1D). As controls, the expression of *Pax7* quickly decreased upon SC activation whereas *MyoD* level sharply increased and peaked at 48 hr. The expression dynamics of *Myc, Rora, Pknox2, Sox4* largely recapitulated the RNA-seq data whereas *Bcl6* and *Esr1* levels exhibited continuous elevation during the entire course (Figure 1D). At the protein level by Western blot, consistent dynamics were observed with the exception of Rora protein, which was not detected in FISCs despite the high RNA level (Figure 1E).

Of note, recent study demonstrated that the isolation process can induce SC activation(51). To circumvent the issue, *in situ* fixation with paraformaldehyde (PFA) prior to isolation was shown to prevent the activation allowing extraction of SCs possibly close to quiescent stage. To pinpoint the expression of the above key TFs in early stages, we examined their expressions using published RNA-seq from SCs fixed by PFA prior to isolation (QSC-T0) and FISCs purified by the standard 3 or 5-hour-long protocol(51) (FISC-T3 and FISC-T5) (Supplementary Figure S2A). Interestingly, the expressions of *Sox4* and *Myc,* particularly *Myc,* were found to be induced in FISCs vs. QSCs, whereas the other four TFs decreased. To validate the above findings, we also isolated RNAs from PFA-fixed QSCs and FISCs and examined their expressions by qRT-PCR (Supplementary Figure S2B and C), we noticed decreased expression of *Pax7* and induced expression for *Atf3/c-Jun*(51); the expressions of the above six TFs, however, all showed evident decrease. We reasoned that the expression levels of these key TFs are very sensitive to the isolation process; the discrepancy between RNA-seq and our qRT-PCR may be caused by difference in isolation duration (our isolation process took about 10 hrs).

Next, to test the functionality of these key TFs, we employed loss-of-function assays by siRNAs in cultured SCs. To this end, FISCs from Pax7-nGFP mice were cultured for two hrs followed by transfection of siRNAs against each TF (Figure 1F); successful knockdown was confirmed (Supplementary Figure S2D). Since it is believed that it takes SCs about 24-36 hrs to enter the first cell cycle and become activated(52), the degree of SC activation was measured by EdU incorporation assay 24 hrs after seeding (Figure 1F). As shown in Figure 1F-G, knockdown of Myc, Bcl6 or Pknox2 led to reduced EdU incorporation thus the delay of SC activation, which however was not observed in the cases of Rora, Sox4 or Esr1 knockdown. To strengthen the above findings, we transiently overexpressed these TFs in cultured SCs (Supplementary Figure S2E). Unexpectedly, forced expression of these TFs failed to result in any obvious changes in the rate of SC activation (Supplementary Figure S2F). Together, these data suggested that Myc, Bcl6, Pknox2 may potentially play a role in SC activation. Nevertheless, the *ex vivo* system prevents the accurate analysis of SC quiescence and early activation stages. We thus reasoned it is imperative to establish an *in vivo* platform to manipulate TF expression in endogenous QSCs without the need to isolate them, this will allow screening of TFs or any factors key to regulate early steps of SC lineage progression.

### Generation and characterization of muscle-specific Cas9 expressing mice

As it is time consuming and expensive to generate knockout mice for each TF using traditional approach relying on the generation of genetically modified mice by transgenesis or gene targeting in embryonic stem cells, we sought to leverage a Cre-dependent Rosa26-Cas9 knockin transgenic mouse line that allows for tissue-specific expression of Cas9. Combined with the relative ease to deliver sgRNAs postnatally, it has become a highly useful tool to study gene function *in vivo*(39,41,53–55). To establish a versatile *in vivo* platform that allows muscle restricted genome editing, we crossed homozygous Cre dependent Rosa26_Cas9-EGFP_ knockin mouse with a Pax7_Cre_ mouse and the resulting progenies were referred to as Pax7_Cas9_ mice (Figure 2A). Co-expression of an enhanced green fluorescent protein (EGFP) allows us to monitor the Cas9-expressing cells. To validate the expression of Cas9 protein in SC, SCs were isolated from six-week-old Pax7_Cas9_ mouse based on GFP expression. Considering the Pax7_Cre+/-_ control mouse does not permit sorting based on GFP expression, Pax7-nGFP mouse was used as sorting control (Ctrl). As expected, two distinguishable sub-populations appeared on FACS sorting plot based on GFP expression (Figure 2B), which is analogous to the Ctrl plot (Supplementary Figure S1A). High expression of Cas9 mRNA or protein was detected in GFP+ cells from the Pax7_Cas9_ mice (Figure 2C and D) but not SCs from the Ctrl mice, confirming the successful expression of Cas9 in SCs. In addition, by Western blotting, Cas9 protein was found to be highly expressed in muscle tissue but not other parts (heart, liver, spleen, lung, kidney and brain) of Pax7_Cas9_ mice or in the skeletal muscle of Pax7_Cre+/-_ Ctrl mice (Figure 2E). Since Pax7 expression emerged as early as embryonic day 10.5 (E10.5) in the muscle progenitor cells to drive the formation of muscle compartments(56), the detected Cas9 expression in muscle tissue may stem from its expression in not only SCs but also the muscle fibers.

**Figure 2.**
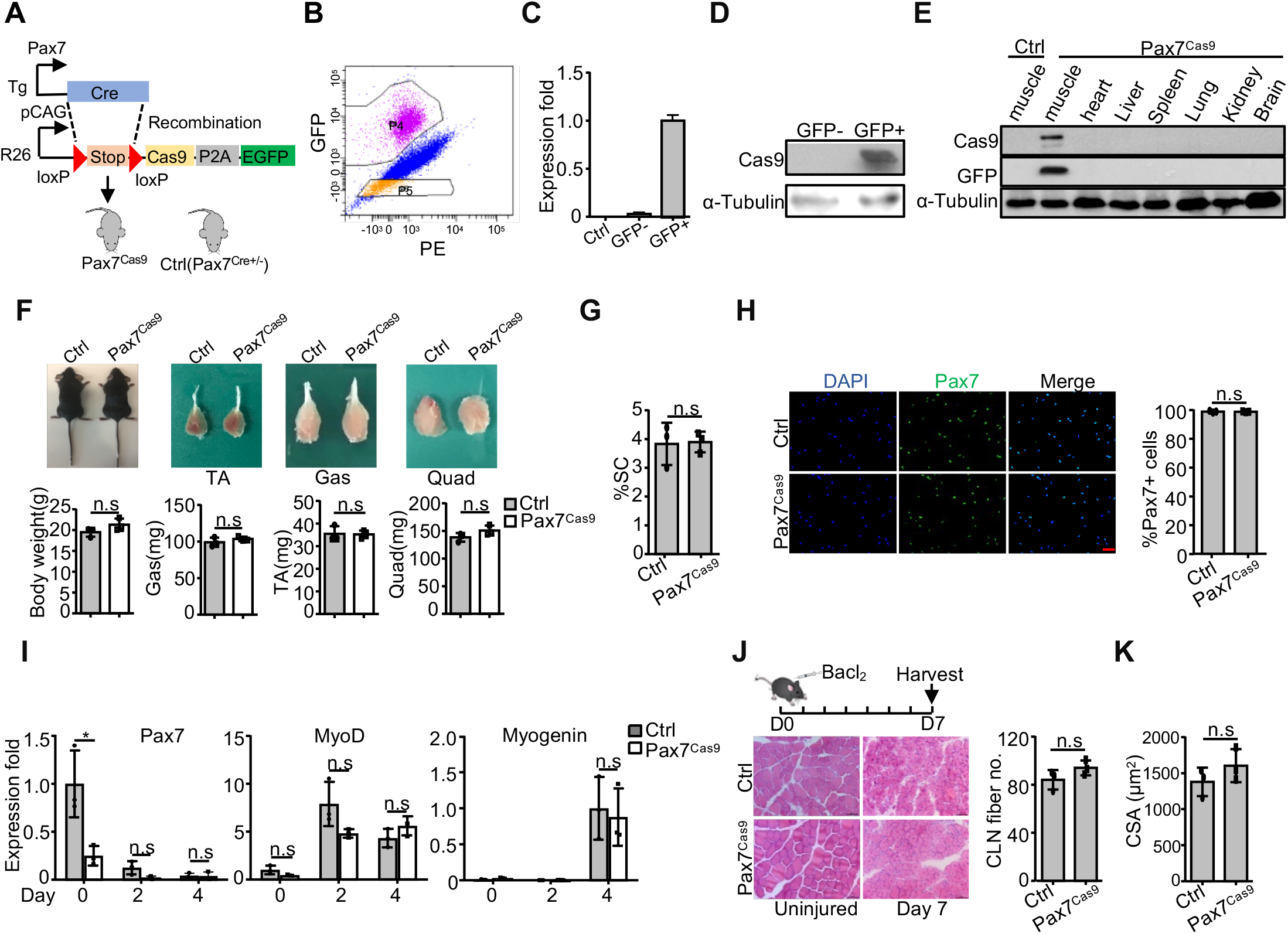
Generation and characterization of muscle-specific Cas9 expressing mice. (**A**) Schematic illustration to show the generation of muscle-specific Cas9 knockin mouse (Pax7_Cas9_). Tg: transgenic; Stop: 3×polyA signal; R26: Rosa26 locus; pCAG: CAG promoter. (**B**) FACS plot showing the gating strategy to isolate Cas9-GFP positive SCs from the Pax7_Cas9_ mouse. P4 represents GFP+ SCs and P5 represents GFP-cells. (**C**) Relative expression of Cas9 mRNAs in the above harvested GFP- and GFP+ cells were detected by qRT-PCR. Cells from Pax7-nGFP mouse were used as Ctrl. (**D**) The expression of Cas9 protein in the above GFP+ or GFP-cells was examined by Western blot. α-Tubulin was used as a loading control. (**E**) The expressions of Cas9 and GFP proteins in the indicated organs of the Pax7_Cas9_ mouse were examined by Western blot. The skeletal muscle from the Pax7_Cre+/-_ mouse was used as control. α-Tubulin was used as a loading control. (**F**) No obvious difference was detected comparing the body mass and three major types of limb muscle (TA: Tibialis anterior; Gas: Gastrocnemius; Quad: Quadriceps) between control (Pax7_Cre+/-_) and Pax7_Cas9_ littermates (n = 3 in each group). Representative images of mice and muscles are shown in the top. (**G**) The percentage of freshly sorted SC showed no difference between control (Pax7-nGFP) and the Pax7_Cas9_ mice (n = 3 in each group). (**H**) IF staining to examine Pax7 protein expression in SCs purified from control (Pax7-nGFP) /Pax7_Cas9_ mice. The above FISCs were seeded for 24 hrs and stained for Pax7 by IF. The percentage of Pax7 positive cells was quantified and the bar graph represents the average of three independent experiments ± s.d. Scale bar, 50 μm. (**I**) The above SCs were cultured for two or four days and mRNA expressions of *Pax7, MyoD* and *Myogenin* were examined (n = 3 in each group). (**J**) TA muscles from eight weeks old control (Pax7_Cre+/-_)/Pax7_Cas9_ mice were injected with BaCl_2_ and subject to H&E staining on day 7 post injury. The regenerating myofibers with centrally localized nuclei (CLN) per field were quantified from three pairs of mice (Right). Scale bar, 100 μm. (**K**) The cross-sectional area (CSA) of the newly formed fibers with CLN was quantitated (n = 3 in each group). All qRT-PCR data were normalized to 18S or GAPDH mRNA and presented as mean ± s.d. *P<0.05. ns, no significance.

To examine any potential effects of Cas9 expression on mice, we found that in line with previous studies(41,53), the resulting Pax7_Cas9_ progenies were born at a Mendelian frequency and appeared viable, healthy and fertile, showing no overt morphological abnormalities; the body size and weight, as well as the limb muscles all appeared comparable to the Pax7_Cre+/-_ Ctrl mice (Figure 2F). To further evaluate whether prolonged expression of Cas9 has any impact on SC homeostasis, we found that a similar percentage of SC were sorted by FACS from Pax7_Cas9_ and Pax7-nGFP Ctrl mice (Figure 2G), indicating Cas9 expression did not affect the number of SCs in muscle. Furthermore, when the sorted cells were then cultured to activate and differentiate; the lineage progression did not appear to be impacted by Cas9 expression as comparable expressions of *MyoD* and *Myogenin* were detected in Pax7_Cas9_ and Ctrl cells (Figure 2I). Although a decreased level of *Pax7* mRNA was detected in FISCs from Pax7_Cas9_ cells but Pax7 protein level measured by immunofluorescence (IF) staining displayed no obvious difference (Figure 2H). Lastly, to further investigate the effect of ectopic expression of Cas9 on SC function, acute muscle injury was induced by injection of barium chloride (BaCl_2_) into the TA muscles of Pax7_Cas9_ or Ctrl (Pax7_Cre+/-_) mice. Hematoxylin and eosin (H&E) staining 7 days after injury revealed no difference in the degree of regeneration as measured by the number of regenerating myofibers featured by centrally localized nuclei (CLN) (Figure 2J) and the cross-sectional areas (CSA) of the newly formed fibers (Figure 2K). Altogether, the above results demonstrated that Cas9 expression in SCs did not impact SC homeostasis and regenerative capacity, warranting the application of the Pax7_Cas9_ mouse for the *in vivo* editing of genes of interest. Of note, we should point out that the above used mice were all heterogeneous Cas9 mice. In some homozygous mice, we found Cas9 was expressed in multiple tissues possibly because Pax7 was expressed in a rare subpopulation of spermatogonia of mice(57) thus turned on Cas9 expression as early as in the zygote. In addition, the number of FISCs from the homozygous mice was significantly reduced compared to heterogeneous or Ctrl mice, showing excessive expression of Cas9 or EGFP may have impact on SC homeostasis(58).

### AAV9-sgRNA mediated editing of *MyoD* in SCs of Pax7_Cas9_ mice

After obtaining the above mouse expressing Cas9 in SCs, we sought to test if it can be used for gene editing of the identified TFs in SCs *in vivo.* As a proof-of-concept, the *MyoD* gene was selected due to the wealth of knowledge of its functions in SC as a master TF orchestrating myogenic program(10,11,17). Prior studies have shown that although disruption of MyoD has no obvious deleterious muscle defect during embryogenesis and postnatal development, its deficiency causes abnormal elevation of SC number on freshly isolated fibers(10). Inducible knockout of *MyoD* in adult mice also results in SC accumulation in the damaged region without fusion after injury(11). Previously, we have also successfully edited *MyoD* in C2C12 myoblasts using CRISPR/Cas9 tool, which led to failure of myoblast differentiation(17). This time a total of six sgRNAs against the first coding exon of *MyoD* were designed by the online tool Crispor(31) (http://crispor.tefor.net/) to minimize potential off-target effect (Figure 3A). Each sgRNA was cloned into a Cas9-EGFP expressing vector and transfected into C2C12 cells followed by FACS based sorting to enrich transfected cells (Figure 3B). Surveyor nuclease assay(30), which enables calculation of the frequency of insertion/deletion (indel) mutations caused by CRISPR/Cas9, was used to examine the cleavage efficiency and screen for the most effective sgRNAs. As shown in Figure 3C, sgRNA2 targeting *MyoD* exon 1 showed the highest editing efficiency (19.5%) and was chosen as the functional sgRNA for further use. To deliver the sgRNA *in vivo*, AAV9 virus was employed because it displays tropism to skeletal muscle and allows transgene expression during a long period of time(22,24). The selected sgRNA2 was placed into a pAAV9-sgRNA vector on which its expression is driven by a human U6 promoter (Figure 3D). This AAV9 vector also carries a fluorescent DsRed gene controlled by a CMV promoter to facilitate the evaluation of transduction efficiency. To test the performance of this AAV9-sgMyoD *in vitro*, it was transfected into C2C12 cells together with a Cas9-expressing plasmid, which resulted in an obvious cleavage at *MyoD* locus (Supplementary Figure S3A).

**Figure 3.**
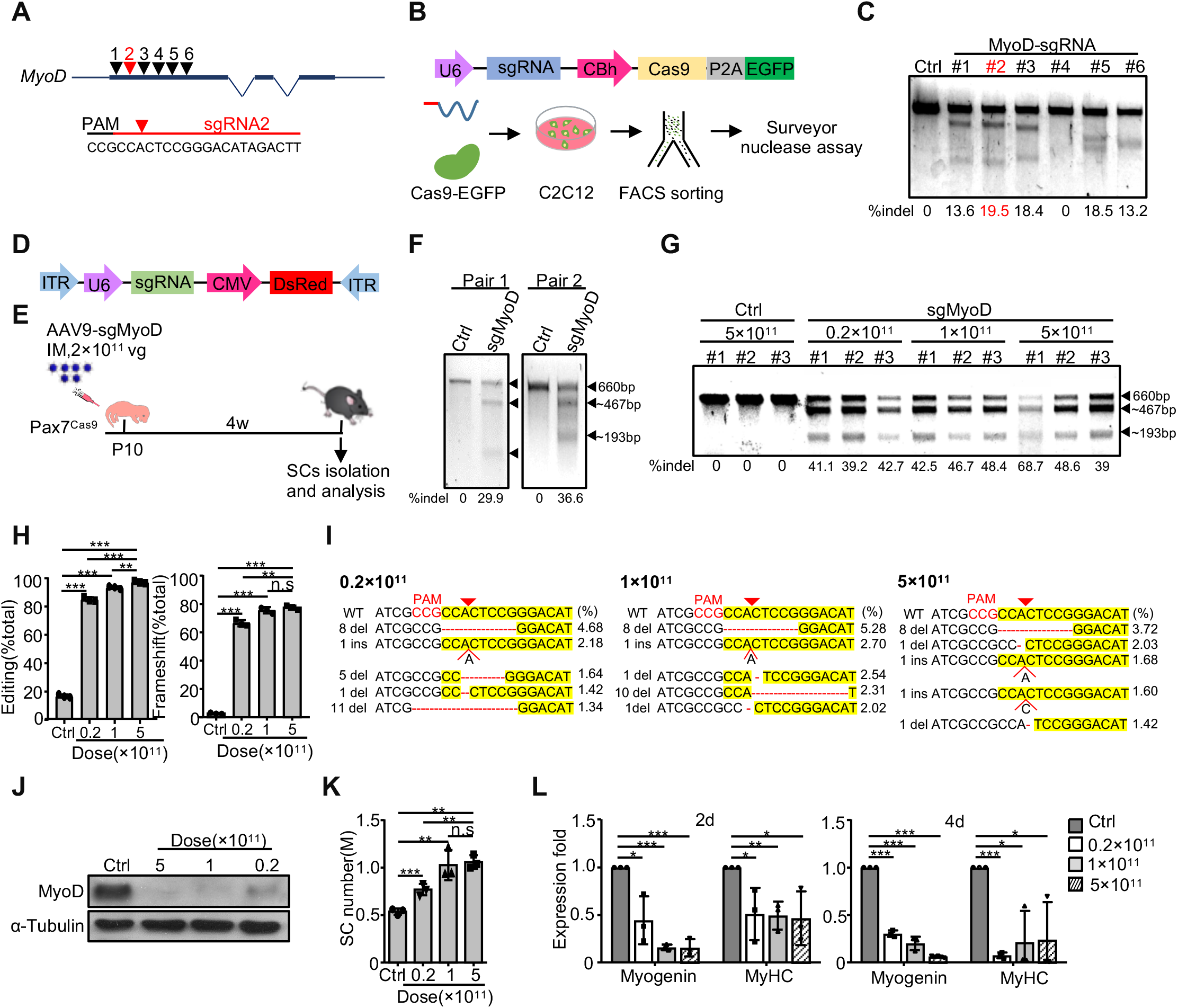
AAV9-sgRNA mediated editing of *MyoD* in SCs of Pax7_Cas9_ mice. (**A**) Design of sgRNAs for targeting the first exon of *MyoD* locus. Selected sgRNA2 was marked as red and its targeted genomic sequence is shown. PAM: protospacer-adjacent motif. (**B**) Top: schematic illustration of the sgRNA-Cas9 expressing vector; Bottom: illustration of *in vitro* sgRNA selection assay. SgRNA-Cas9 expressing plasmid was transfected into C2C12 myoblasts and positively transfected cells were sorted out by GFP expression one day after transfection and cultured for another two days before Surveyor nuclease assay. (**C**) Agarose gel image to show the result of Surveyor nuclease assay on *MyoD* locus. The percentage of indel formation is shown. SgRNA2 with the highest frequency of indel formation is marked as red. (**D**) Schematic illustration of the pAAV9-sgRNA vector for *in vivo* sgRNA expression. (**E**) Schematic illustration of the experimental design for AAV administration and SC isolation and analysis. (**F**) Genomic DNAs from SCs of the Pax7_Cas9_ mice administrated with AAV9-sgMyoD virus were subject to Surveyor assay to test the frequency of indel formation at *MyoD* locus. Wide type (660 bp) and cleaved bands (~467 bp/193 bp) by Surveyor are shown by arrowheads. The percentage of indel formation is shown (n=2 in each group). (**G**) Three doses of AAV virus were administrated and Surveyor assay was performed as above to estimate the indel formation at *MyoD* locus in genomic DNAs from FISCs. Wide type (660 bp) and cleaved bands (~467 bp/193 bp) by Surveyor are shown by arrowheads. (**H**) Quantification of the editing efficiency (Left) and frameshift mutations (Right) determined by deep sequencing of the genomic DNAs from the above (G). The percentage was calculated as the ratio of the edited reads (Left) or frameshift mutations (Right) to the total reads and presented as mean ± s.d (n=3 in each group). (**I**) The five most frequently detected indel classes under each dose of AAV9-sgMyoD virus are shown. The percentage is calculated as the ratio of reads for each indel class to the total edited reads. (**J**) FISCs from the three doses of AAV9-sgMyoD administration were cultured for two days and the MyoD protein level was examined by Western blot. α-Tubulin was used as loading control. (**K**) The number of SCs purified from Pax7_Cas9_ mice administrated with different doses of AAV9-sgMyoD virus is presented as mean ± s.d (n = 3 in each group). M: million. (**L**) The above isolated SCs were cultured for two or four days and the relative expressions of *Myogenin* or *MyHC* was detected by qRT-PCR (n = 3 in each group). All qRT-PCR data were normalized to 18S mRNA and presented as mean ± s.d. *P<0.05, **P<0.01 and ***P<0.001. ns, no significance.

The above AAV9-sgMyoD plasmid was then packed into virus and intramuscularly (IM) injected into the Pax7_Cas9_ mice at postnatal day 10 (P10) with a single dose of 2×10_11_ vg/mouse (Figure 3E). For the control group, the same dose of AAV9 virus containing pAAV9-sgRNA backbone without any sgRNA insertion was injected. We reasoned that active postnatal myogenesis occurs at P10 thus allowing effective genome editing in proliferating myoblasts, which will carry the editing when a sub-population of them return to quiescence to establish the SC pool around 2-4 weeks after birth(59). Four weeks after injection, we first evaluated the transduction efficiency of AAV9 virus into SCs based on DsRed expression. Two distinguishable sub-populations were separated on the plot according to GFP expression as expected (Supplementary Figure S3B), but the GFP+ SCs did not present differential DsRed expression. Further examination of DsRed protein expression in the input cells (Supplementary Figure S3B) led to separation of DsRed-high vs. -low populations and GFP+ cells were located in the DsRed-high peak (Supplementary Figure S3C), indicating most GFP expressing SCs were indeed infected by the virus. Furthermore, above 80% of the isolated SCs were DsRed positive by IF staining (Supplementary Figure S3D), manifesting efficient transduction of the AAV9 virus into SCs. In addition, we noticed that the entire injected muscle was visibly red during dissection, confirming robust infection of AAV-sgRNA virus into the skeletal muscle. To estimate the editing efficiency, genomic DNA from FISCs was subject to Surveyor assay; around 30%~40% indel occurrence was detected in the AAV9-sgMyoD injected mice but not the Ctrl group (Figure 3F), confirming the success in editing *MyoD.* Further analysis of MyoD transcripts from SCs cultured for four days using Surveyor assay also confirmed efficient mutagenesis (Supplementary Figure S3E). Taken together, the above results demonstrated the Pax7_Cas9_/AAV9-sgRNA system can be used for effective gene editing in SCs *in vivo.*

Since a recent study implied that high dose administration of AAV9 virus to nonhuman primates and piglets causes severe toxicity(60), we next sought to determine the minimal dose of AAV9 needed to achieve effective editing. To this end, high (5×10_11_ vg/mouse), middle (1 × 10_11_ vg/mouse) or low (0.2×10_11_ vg/mouse) dose of AAV9-sgMyoD virus particles were intramuscularly injected at P10. Pax7_Cas9_ mice administrated with a high (5×10_11_ vg/mouse) dose of AAV9 virus containing pAAV9-sgRNA backbone without any sgRNA insertion were used as Ctrl. The mice were monitored daily and no obvious abnormalities were observed even in the high dose injected group. Four weeks after injection, GFP+ SCs were sorted out to detect indel formation at *MyoD* locus. Interestingly, the editing efficiency determined by Surveyor assay in FISCs displayed no obvious difference among the three doses (Figure 3G and Supplementary Figure S3F); and similar indel occurrences were also detected in MyoD transcripts from ASCs cultured for two days (Supplementary Figure S3G). To precisely determine the level of editing, sgMyoD targeted locus was amplified from genomic DNA in FISCs and subject to deep sequencing. The analysis revealed that the high dose treatment led to a significantly higher level of mutations than the low (95.08% ± 0.80% vs. 83.16% ± 1.42%) and middle dose groups (91.28% ± 0.92%) (Figure 3H); however, the percentage of frameshift mutations between high and middle dose groups presented no obvious difference (77.43% ± 1.42% vs. 75.58% ± 2.26%) (Figure 3H). The indels with the highest frequency included small deletions (within 10 bp) and +1 bp insertion at target site and the patterns were similar among different doses (Figure 3I), which is in agreement with a prior report showing that the repair outcomes of CRIPSR/Cas9 mediated genome editing are not random(61).

As the consequence of the above editing, we found that MyoD protein was completely eliminated in SCs from mice receiving high or middle dose of the AAV9-sgMyoD virus whereas noticeable MyoD protein in low dose injected group (Figure 3J). This was further confirmed by IF staining which indicated that administration of the virus resulted in clear repression of MyoD protein level in SCs, even for the low dose group (Supplementary Figure S3H and I). Unexpectedly, we failed to observe a sharp decrease of MyoD transcript level (Supplementary Figure S3J). The discrepancy in MyoD mRNA and protein expression indicated that CRISPR/Cas9 mediated genome editing does not necessarily impact mRNA level as previously described(41). In sum, these findings confirmed the high efficiency of this system to knockout MyoD gene expression in SCs *in vivo.* To examine the impact of MyoD loss on SCs, we noticed a robust increase of the number of FISCs (Figure 3K and Supplementary Figure S3K), similar to previous reports using *MyoD* knockout mice(10,11). The middle dose, 1× 10_11_ vg/mouse, yielded a much higher increase than the low dose but increase of the dose to 5 × 10_11_ vg was unable to elevate the number further. Phenotypically, when the cells were cultured for differentiation for two or four days, decreased level of *Myogenin* or *MyHC* expression was found in the mutant vs. Ctrl group (Figure 3L), and the decrease was comparable for the three doses, confirming a defect in myoblast differentiation caused by MyoD loss. Taken together, we decided to use the middle dose (1 × 10_11_ vg/mouse) of AAV9 virus for the subsequent investigation.

### Development of an AAV9-dual sgRNA strategy for precise genome editing in SCs

Although the application of single sgRNA exhibited high efficiency in editing *MyoD* locus, several studies have demonstrated that simultaneous use of dual sgRNAs could induce indel as well as large deletion at target loci to further enhance editing efficiency(39,41,62). We thus sought to develop an AAV9-dual sgRNA system expressing two sgRNAs from a single vector which was shown to display higher cleavage potential compared to sgRNAs from two separate vectors(63) (Figure 4A). Using *MyoD* as a paradigm, two sgRNAs (sgRNA2 and sgRNAc) were selected to remove a 298 bp in exon 1 upon precise ligation of the two predicated cutting sites (Figure 4A and Supplementary Figure S4A). A middle dose of AAV9-dual sgMyoD virus was injected into Pax7_Cas9_ mice to test the editing efficiency *in vivo* (Figure 4B). For comparison, the same dose of single AAV9-MyoD-sgRNA2 virus was administrated. Expectedly, the AAV9-dual sgMyoD virus caused excision of the intervening sequence between the two sgRNA targeting sites (Figure 4C), which was not detected in the single sgRNA or control virus infected SCs. However, the level of MyoD protein showed no difference between the dual and single sgRNA groups (Figure 4D). Consistenly, the AAV9-sgMyoD viruses led to an increase of the FISC number and the dual sgRNAs did not induce further elevation (Figure 4E); the expression of *Myogenin* was also comparable in both groups (Figure 4F).

**Figure 4.**
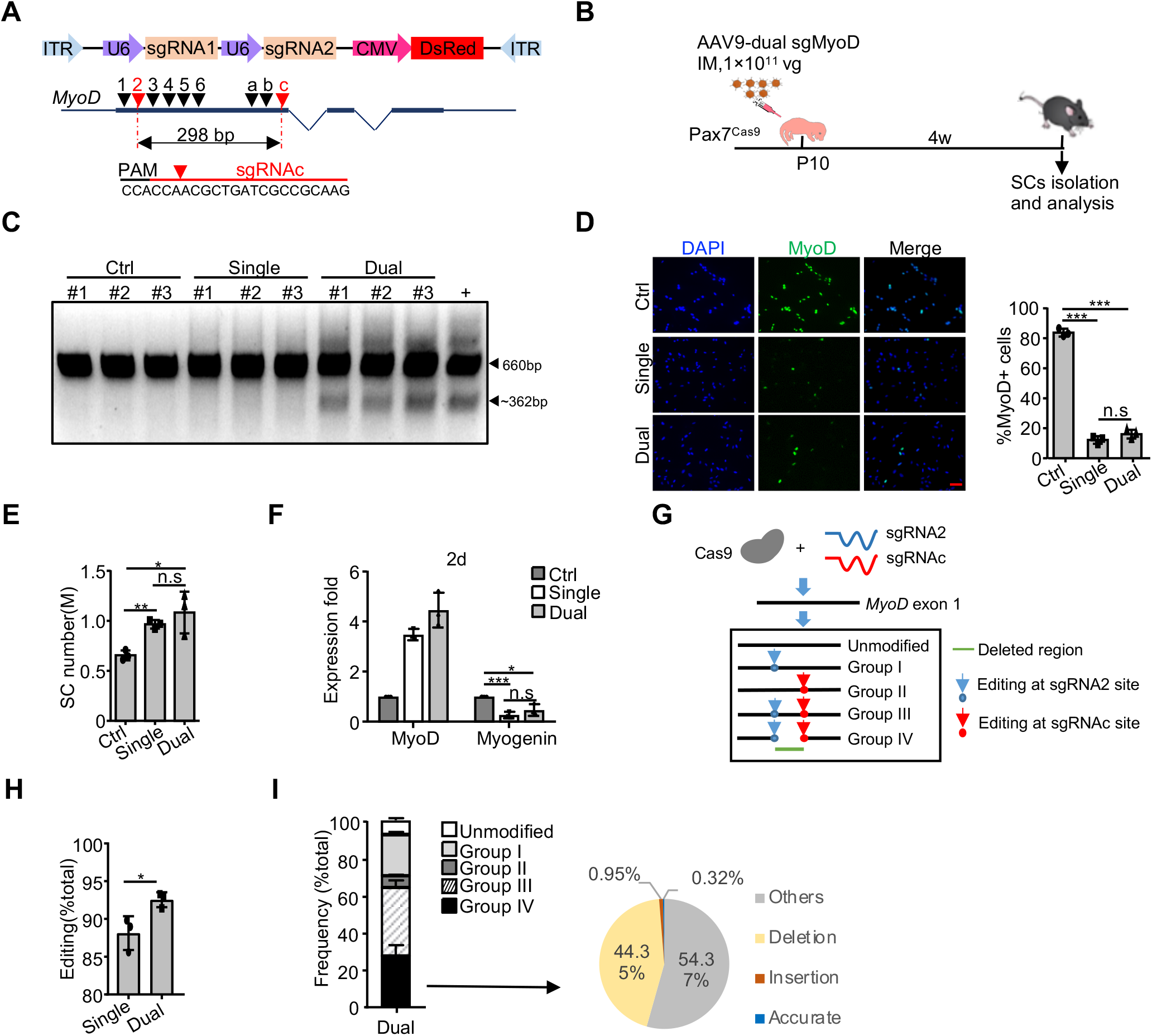
Development of an AAV9-dual sgRNA strategy for precise genome editing in SCs. (**A**) Top: schematic illustration of the pAAV9-sgRNA vector used for dual sgRNAs expression; Middle: design and selection of sgRNAs targeting the first coding exon of *MyoD.* SgRNA2 and sgRNAc selected for *in vivo* use were marked as red. Bottom: the targeted genomic sequence for sgRNAc. PAM: protospacer-adjacent motif. (**B**) Schematic illustration of the experimental design for AAV-dual sgMyoD virus administration and SC isolation/analysis. The administration of middle dose of AAV9-sgRNA backbone (without sgRNA insertion) virus or single AAV9-sgMyoD virus was used as control or comparison. (**C**) PCR analysis to test the cleavage efficiency at *MyoD* locus in FISCs from the above injected mice. DNA from C2C12 cells co-transfected with Cas9-expressing plasmid (pX458) and AAV9-dual sgMyoD vector was used as positive control. Wide type (660 bp) and cleaved bands (~362 bp) are shown by arrowheads. (**D**) SCs were cultured for two days and IF stained for MyoD (Left). The number of positively stained cells was counted (Right) (n = 3 in each group). Scale bar, 50 μm. (**E**) The number of FISCs sorted from the above mice (n = 3 in each group). M: million. (**F**) Relative expression of *MyoD* and *Myogenin* in the above SCs cultured for two days were determined by qRT-PCR. The qRT-PCR data were normalized to 18S mRNA and presented as mean ±s.d (n = 3 in each group). (**G**) Schematic illustration to show non-homologous end joining (NHEJ) repair of double-strand breaks (DSBs) induced by AAV9-dual sgMyoD. The edited products were divided into four groups based on whether the indels formed at either sgRNA or both target sites. Group I: editing at sgRNA2 site only; II: sgRNAc site only; III: individually at sgRNA2 and sgRNAc site; IV: simultaneously at both sites to cause deletion. (**H**) Genomic DNAs were isolated from FISCs and subject to deep sequencing to assess the editing events. The percentage of editing was calculated as the ratio of the edited reads to the total reads (n = 3 in each group). (**I**) Left: distribution of the total sequencing reads in the dual sgRNAs group. Right: distribution of the above Group IV reads. Accurate: reads with precise ligation of the two predicted cutting sites. Insertion (or deletion): reads with insertion (or deletion) at the junction site. Others: reads that do not belong to the above. The percentage was calculated as the ratio of each part to the total reads in group IV (n = 3 in each group). All the bar graphs are presented as mean ± s.d. *P<0.05, **P<0.01 and ***P<0.001. ns, no significance.

To further compare the frequency of indel formation between the single and dual sgRNAs treatment, deep sequencing was performed on genomic DNA from FISCs. The edited products for dual sgRNA2/c were categorized into four groups, I-IV, based on whether the indels formed at sgRNA2 site only, sgRNAc site only, individually at both sites (without deleting target locus) or simultaneously at both sites (deletion of target locus) (Figure 4G). In total, the dual sgRNAs resulted in above 90% editing at *MyoD* locus, which was slightly higher than the single sgRNA group (92.55% ± 0.99% vs. 88.13 ± 2.23%) (Figure 4H). For the dual-sgRNA group, sgRNA2 and sgRNAc presented 86.2% and 70.21% indel formation respectively (Supplementary Figure S4B). To further examine if the deletion was precise ligation of the two predicted cutting sites as in the case of CRISPR/Cas9-mediated short time deletion (within days)(40,63), we next closely examined the dual sgRNAs induced deletion. About 27% of the total reads were detected as deletion (Group IV, Figure 4I), among which only 0.32% was yielded from the precise ligation. Recent study also indicated that Cas9 cleaves the non-complementary DNA strand with a flexible profile to produce not only blunt ends but also 5’ overhanging ends with 1-3 nucleotides, which contributes to the formation of 1-3 insertions at the junction of two cutting sites(40). However, in our case, the insertions only occurred in 0.95% of the deleted reads and about half was deletion around the junction (Figure 4I). We thus examined the target sequences of these two sgRNAs closely and realized a new sgRNA targeting site would form as a result of precise ligation of the two cutting ends, and coincidently this new site could be further edited by sgRNAc (Supplementary Figure S4C). Therefore, the outcomes of CRISPR/Cas9-mediated long time editing (within months) need further investigation. Together, our findings demonstrated the dual sgRNAs editing system yielded higher editing efficiency than single sgRNA thus was chosen for subsequent genome editing of the identified key TFs.

### Attempts of CRISPR/Cas9/AAV9 mediated genome editing in quiescent SCs

In the described strategies above, AAV-sgRNA administration into whole muscle leads to a caveat that in addition to SCs, myofibers will also be edited. Moreover, editing at P10 stage may affect SC pool establishment and complicate the subsequent study of SC function in adult regeneration. To test if it is possible to edit QSCs only after postnatal myogenesis ceases, we attempted the *in vivo* genome editing using an inducible Cas9 expressing mouse by crossing the homozygous Rosa26_Cas9-EGFP_ knockin mouse with the Pax7_CreER_ mouse; the resulting offsprings were referred to as Pax7_ER-Cas9_ mice (Figure 5A). As the Cre recombinase only functions in Pax7 positive cells after exposure to Tamoxifen (Tmx)(64), Cas9 induction can be controlled by Tmx injection in the resulting progenies. High dose of single AAV9-sgMyoD virus was injected through IM into four weeks old Pax7_ER-Cas9_ mice followed by five consecutive intraperitoneal (IP) administrations of Tmx to induce Cas9 expression two weeks later. The mice were sacrificed for SC isolation after another three weeks (Figure 5B). Two distinguished sub-populations were captured by FACS sorting based on GFP expression, demonstrating sufficient induction of Cas9 in SCs after Tmx injection (Figure 5C). Despite previous study shows QSCs are refractory to AAV9 transduction in adult mice(25), successful AAV9 transduction in SCs was confirmed by PCR amplification of DsRed specifically in DNAs from AAV9 infected FISCs but not control Pax7_ER-Cas9_ mice without AAV9 administration, whereas PCR product targeting Cas9 coding region was detected in all the tested mice (Figure 5D). Despite the above successful AAV9 transduction into SCs, no editing at *MyoD* locus (Figure 5E) or change of MyoD, Myogenin, or MyHC expression (Figure 5F) was detected in the AAV9-sgMyoD virus injected group. Next, we performed the injection of the AAV9-sgMyoD virus at P10 hoping the increased duration would further improve the transduction efficiency(22,24). The actively proliferating myoblasts were expected be infected by the virus and a portion of them would return to quiescence carrying the sgRNAs which allows the editing to occur after Cas9 expression is induced with the Tmx injection (Figure 5G). Compared to the first strategy, clear shifting of SCs to the DsRed positive population was noticed during FACS based isolation (Figure 5H) and PCR analysis of DsRed coding region in DNAs from FISCs also indicated successful AAV9 transduction (Figure 5I). Still, no genome editing occurred at *MyoD* locus as examined by Surveyor nuclease assay (Figure 5J); consistently, the MyoD protein level did not exhibit obvious change (Figure 5K).

**Figure 5.**
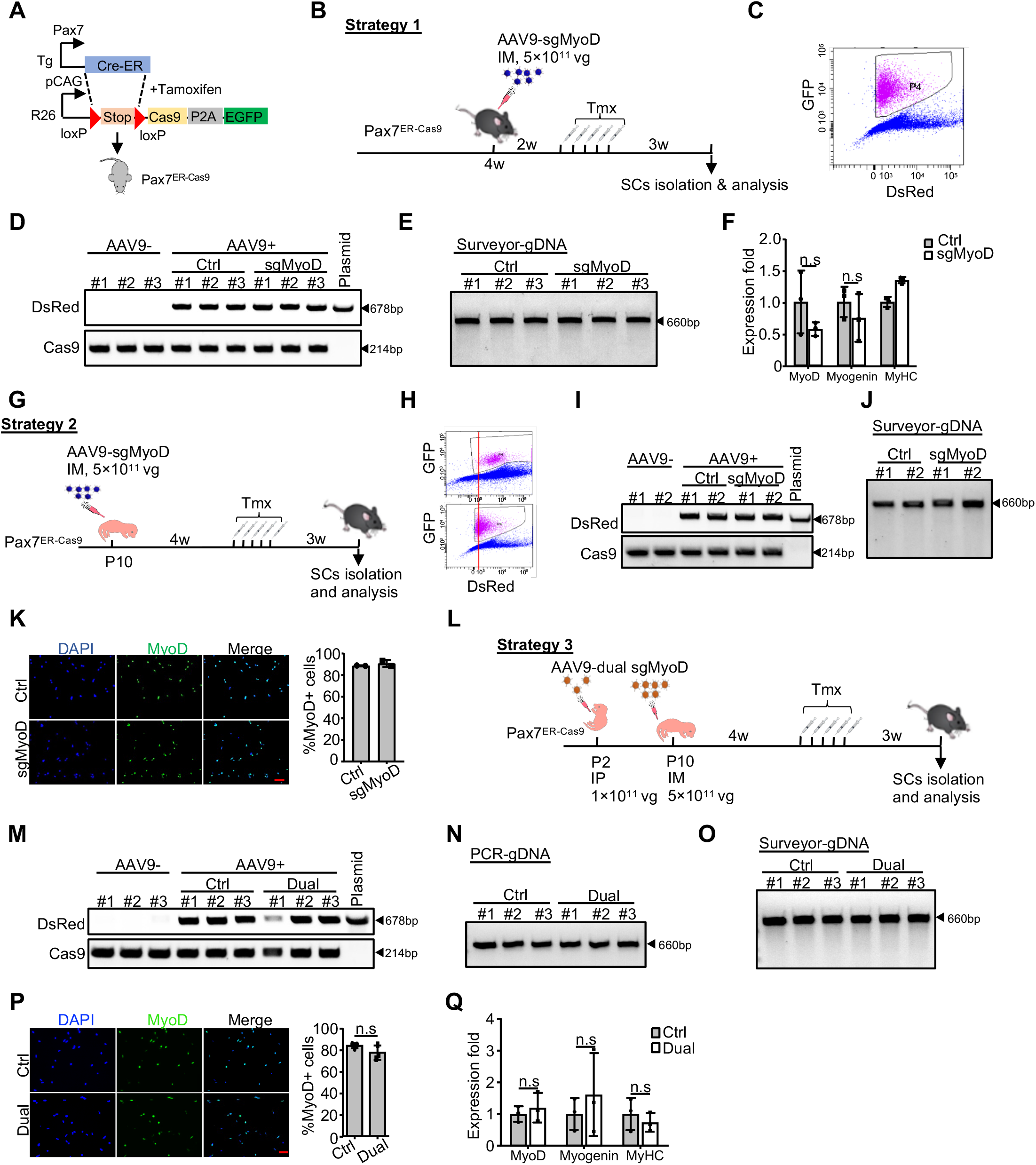
Attempts of CRISPR/Cas9/AAV9 mediated genome editing in quiescent SCs. (**A**) Schematic illustration to show the generation of the inducible muscle-specific Cas9 knockin mouse (Pax7_ER-Cas9_). (**B**) Schematic illustration of the first strategy to edit quiescent SCs (QSCs) in the above generated mouse. High dose of AAV9-sgMyoD or control virus was injected through IM to four weeks old Pax7_ER-Cas9_ mice. After two weeks, Tamoxifen (Tmx) was injected intraperitoneally (IP) for five consecutive days to induce Cas9 and GFP expression. The mice were sacrificed for SC isolation and analysis after another three weeks. (**C**) FACS plot showing efficient induction of Cas9-GFP expression in SCs from the above mice. The gate P4 indicates the population of Cas9-GFP positive SCs isolated for following analysis. (**D**) Genomic DNAs (gDNA) were isolated from the above sorted SCs and PCR was performed to amplify the DsRed or Cas9 coding region. Genomic DNA from Pax7_ER-Cas9_ mice without AAV9 administration was used as negative control and the AAV9-MyoD-sgRNA2 plasmid was used as positive control. (**E**) Surveyor assay was performed to detect the editing efficiency at *MyoD* locus in the above FISCs. A 660 bp amplicon of wild type MyoD locus is shown. (**F**) Relative expression of *MyoD, Myogenin* and *MyHC* mRNAs were detected in the above SCs cultured for four days (n = 3 in each group). (**G**) Schematic illustration of the second strategy to edit QSCs in Pax7_ER-Cas9_ mice. High dose of AAV9-sgMyoD or control virus was injected through IM at P10. Four weeks later, Tmx was injected via IP for five consecutive days and the mice were sacrificed for SC isolation and analysis after another three weeks. (**H**) FACS plot showing the gating strategy to sort out Cas9-GFP positive SCs from the above treated mice (Upper panel). A stronger DsRed signal compared to Strategy 1 (Lower panel) indicates a higher transduction efficiency. (**I**) Transduction of AAV9 virus in SCs was examined as describe in (D). (**J**) Editing efficiency was examined by Surveyor assay as in (E). (**K**) SCs from the above treated mice were cultured for two days and IF stained for MyoD (Left) and the number of positively stained cells was counted (Right) (n=2 in each group). Scale bar, 50μm. (**L**) Schematic illustration of the third strategy to edit QSCs from adult mice. A middle dose of AAV9-dual sgMyoD or control virus was injected through IP into Pax7_ER-Cas9_ mice at P2 and another high dose of the same virus was administrated at P10 through IM. Four weeks later, Tmx was administrated via IP for five consecutive days and the mice were sacrificed for SC isolation and analysis after another three weeks. (**M**) Transduction of AAV9 virus in SCs was examined as describe in (D). (**N-O**) Editing efficiency was examined by PCR analysis targeting *MyoD* locus (**N**) or Surveyor assay(**O**) as in (E). (**P**) SCs from the above treated mice were cultured for two days and IF stained for MyoD (Left) and the number of positively stained cells was counted (Right) (n = 3 in each group). Scale bar, 50μm. (**Q**) Relative expressions of *MyoD, Myogenin* and *MyHC* mRNAs were detected in the above SCs cultured for four days (n = 3 in each group). All qRT-PCR data were normalized to 18S or GAPDH mRNA. All the bar graphs are presented as mean ± s.d. ns, no significance.

Since dual sgRNAs led to a better performance than single sgRNA (Figure 4), we further applied the AAV9-dual sgMyoD virus. In addition to an IM administration at P10, we added an extra administration at P2 aiming to improve the transduction efficiency (Figure 5L); successful AAV9 transduction was detected (Figure 5M), which however led to unsuccessful editing (Figure 5N-Q). Taken together, our results demonstrated that the *in vivo* CRISPR/Cas9/AAV9 system failed to induce efficient genome editing at the *MyoD* locus in QSCs.

### CRISPR/Cas9/AAV9 mediated genome editing of *Myc* hindered SC activation and muscle regeneration

After the above in-depth evaluation of the performance of the CRISPR/Cas9/AAV9 mediated genome editing at *MyoD* locus, we next employed this system to study the key TF functions in SC quiescence maintenance and activation. *Myc* was a preferable target considering its known role in inhibiting myoblast differentiation(45) and the potential function showing in Figure 1G from using siRNA oligos *ex vivo* in SCs. Two sgRNAs (sgRNA1 and sgRNAb) which would induce frameshift of *Myc* coding region were selected through Surveyor nuclease assay in C2C12 cells (Figure 6A and Supplementary Figure S5A). AAV9-dual sgMyc succeeded in excising the *Myc* locus *in vitro* as expected (Supplementary Figure S5B). Following the above established administration strategy (Figure 6B), a dose of 5×10_11_ vg/mouse of the dual-sgRNA virus was injected to the Pax7_Cas9_ muscle at P10. PCR analysis using genomic DNAs from FISCs (Figure 6C) and Myc transcripts (Supplementary Figure S5C) revealed *Myc* locus was successfully edited resulting in nearly 235 bp deletion, demonstrating efficient editing. When using the virus dose of 1×10_11_ vg, which, however, led to insufficient editing (Supplementary Figure S5D and E), showing an AAV dose-dependent editing. Subsequent analysis revealed that although the level of Myc transcripts was not significantly changed (Figure 6D), Myc protein level was evidently decreased (Figure 6E). Consistent to the finding from *MyoD* locus, no indel occurrence was detected when attempting the editing in QSCs (Supplementary Figure S5F and G).

**Figure 6.**
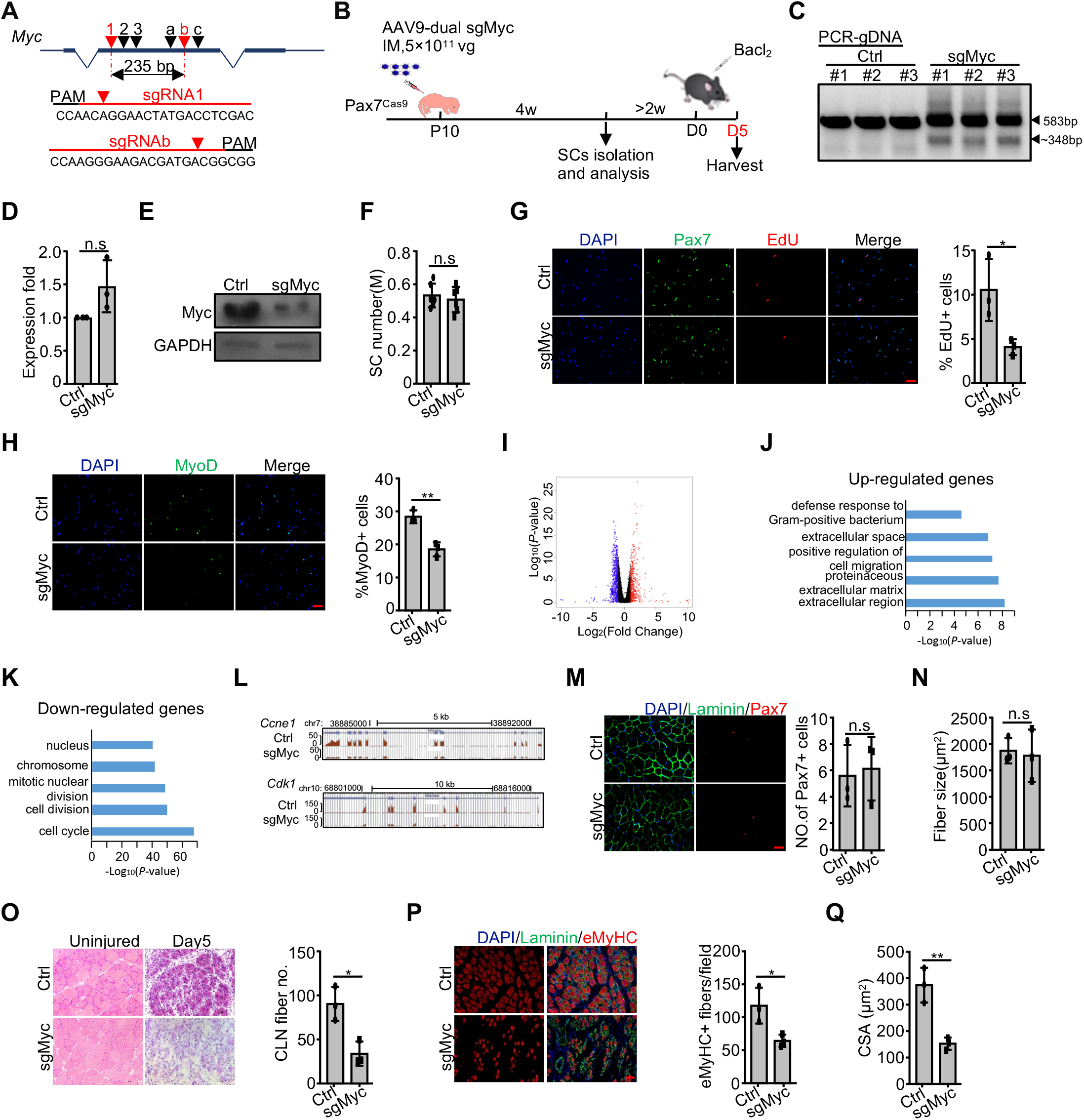
CRISPR/Cas9/AAV9 mediated genome editing of *Myc* hindered SC activation and muscle regeneration. (**A**) Design and selection of sgRNAs targeting *Myc* locus. SgRNA1 and sgRNAb were selected to delete a 235 bp of exon 2. (**B**) Schematic illustration of the experimental design for *in vivo* genome editing of *Myc* locus and analysis of its effect on SCs and muscle regeneration. Pax7_Cas9_ mice were administrated with high dose of control or AAV9-dual sgMyc virus at P10 and SCs were isolated four weeks later for analysis. To assay for muscle regeneration, BaCl_2_ was intramuscularly injected at the TA region of one leg at least six weeks after AAV9 virus administration. The injected muscle was harvested five days post injury and subject to analysis. (**C**) Genomic DNAs were isolated from the above sorted SCs and PCR analysis was performed to test the cleavage efficiency. Wide type (583 bp) and cleaved fragments (~348 bp) are shown by arrowheads. (**D**) The above SCs were cultured for 24 hrs and relative expression of *Myc* mRNAs was detected by qRT-PCR. The qRT-PCR data were normalized to 18S mRNA (n=3 in each group). (**E**) Myc protein level in the above SCs was examined by Western blot. GAPDH was used as loading control. (**F**) The number of SCs isolated from the above control or dual sgMyc virus treated mice (n = 6 in each group). M: million. (**G**) The above FISCs were labeled with EdU for 24 hrs and the percentage of EdU positive cells was quantified (n = 3 in each group). Scale bar, 50 μm. (**H**) The above FISCs were IF stained for MyoD expression four hrs after seeding (Left) and the percentage of positively stained cells was quantified (Right) (n = 3 in each group). Scale bar, 50 μm. (**I**) RNAs were isolated from the above SCs cultured for 24 hrs and subject to RNA-seq analysis. Differentially expressed genes were identified with blue and red dots representing down- or up-regulated genes, respectively. (**J-K**) GO analysis of the above up- or down-regulated genes was performed and the top five enriched items are shown in y axis. X axis shows the -log10[*P* value] of the GO terms. (**L**) Genomic snapshots showing two examples of significantly down-regulated genes in sgMyc vs. ctrl treatment. (**M**) Immunostaining of Pax7 (red) and Laminin (green) waere performed on the uninjured contralateral TA muscle (Left). The number of Pax7 positive cells per 100 fibers was counted (Right) (n = 3 in each group). Scale bar, 50 μm. (**N**) Average fiber size of the above TA muscle was quantified (n = 3 in each group). (**O**) H&E staining was performed 5 days post the BaCl_2_ injection. Uninjured TA muscle was used as control (Left). The regenerating myofibers with CLN per field was quantified (Right) (n = 3 in each group). Scale bar, 100 μm. (**P**) Immunostaining of eMyHC (red) and Laminin (green) on the above TA muscle sections (Left) and the number of eMyHC+ myofibers was counted (Right) (n = 3 in each group). Scale bar, 50 μm. (**Q**) The cross-sectional area (CSA) of the newly formed fibers with CLN was quantified (n = 3 in each group). All the bar graphs are presented as mean ± s.d. *P<0.05, **P<0.01. ns, no significance.

When analyzing the impact of *Myc* editing on SCs, no obvious difference in the number of sorted SCs was detected in AAV9-dual sgMyc vs. Ctrl group (Figure 6F and Supplementary Figure S5H), suggesting the SC homeostasis may not have been altered by *Myc* editing at P10. Similar to the observation in the siRNA knockdown cells (Figure 1G), Myc loss led to a significant delay of SC activation as measured by reduced EdU incorporation 24 hrs in culture (Figure 6G and Supplementary Figure S5I) and decreased number of MyoD+ cells four hrs after isolation (Figure 6H). To gain more insight into how Myc regulates SC early activation, we conducted RNA-seq analysis to globally characterize transcriptomic changes induced by Myc depletion. A total of 519 genes were up-regulated, whereas 840 genes were down-regulated after Myc deletion (Figure 6I and Supplementary Data S7). Subsequent GO analysis revealed that the up-regulated genes were enriched for extracellular region-related terms (Figure 6J and Supplementary Data S7) and the down-regulated genes were enriched for cell cycle-related terms (Figure 6K and Supplementary Data S7), which was consistent with the inhibited SC activation. For example, the expression of *Ccne1* and *Cdk1*, which are essential to G1/S phase transition, were obviously repressed in sgMyc SCs (Figure 6L), indicating that Myc may be involved in SC early activation through regulating cell cycle progression.

To further examine whether loss of Myc impairs damage induced muscle regeneration, Pax7_Cas9_ mice were administrated with high dose of AAV9-dual sgMyc virus at P10 followed by BaCl_2_ in the TA muscle of one side after at least six weeks (Figure 6B). The body weight and muscle weight of the uninjured contralateral limb did not display evident difference compared to littermates injected with control virus (Supplementary Figure S5J). The SC number was not affected in sgMyc vs. control group prior to the injury (Figure 6M), further indicating that Myc may not play a role in regulating homeostasis of the SC pool. And the average fiber size showed no difference compared to the control (Figure 6N). However, five days post-injury, a much lower number of regenerating myofibers with CLN was observed in sgMyc vs. control group (Figure 6O). Consistently, immunostaining of eMyHC, a marker of newly formed fibers, revealed that the number of eMyHC+ fibers was markedly decreased (Figure 6P); measurement of fiber size also revealed a decreased cross-sectional area in sgMyc vs. control group (Figure 6Q). Together, the above findings demonstrated the critical requirement of Myc for proper muscle regeneration after acute injury.

### CRISPR/Cas9/AAV9 mediated genome editing of *Bcl6* and *Pknox2* led to distinct effects on SC activation and muscle regeneration

To further solidify the usefulness of our system in editing TFs and studying their functions *in vivo*, we next applied it to *Bcl6* which appeared to be functional in SCs *ex vivo* showing by siRNA knockdown (Figure 1G). The selected combination of sgRNA2/3 (Figure 7A) targeting exon 3 showed efficient cleavage both *in vitro* and *in vivo* (Figure 7B-C and Supplementary Figure S6A-B), whereas failed to edit the *Bcl6* locus in QSCs (Supplementary Figure S6C and D). Similar to the *Myc* case, administration of high dose of AAV9-dual sgBcl6 virus at P10 did not change the number of sorted FISCs (Figure 7D). When analyzing the cell activation by EdU incorporation assay, dual sgBcl6 cells displayed obvious acceleration compared to Ctrl cells (Figure 7E), hinting the repressive role of Bcl6 during SC activation; this is opposite to the finding from using siRNAs *ex vivo* (Figure 1G). However, no significant change of the number of MyoD+ cells was found (Supplementary Figure S6E), suggesting that Bcl6 inhibits SC activation without affecting MyoD expression; actually previous study has shown that induction of Bcl6 required MyoD expression(46). Next, we further applied RNA-seq in sgBcl6 infected SCs in culture for 24 hrs. A total of 76 down- and 129 up-regulated genes were detected in sgBcl6 vs. Ctrl group (Figure 7F and Supplementary Data S8). The number of down-regulated genes was insufficient for GO analysis; the up-regulated genes were found to be enriched for GO terms such as “extracellular region” and “extracellular matrix” (Figure 7G and Supplementary Data S8). For example, the expressions of two extracellular matrix (ECM) related genes, *Lcn2* and *Mmp3,* were obviously up-regulated in sgBcl6 vs. Ctrl cells (Figure 7H). It is known that remodeling of ECM including collagen and proteoglycans is essential to SC function and muscle regeneration(65). *Lcn2* which is an ECM regulator modulates SC activation through its association with the ECM protease matrix metalloproteinase-9 (Mmp-9)(66) and the Mmps have been reported to be involved in mechanical stretch-induced SC activation(67). Thus, Bcl6 may coordinate SC activation through regulating these ECM related genes.

**Figure 7.**
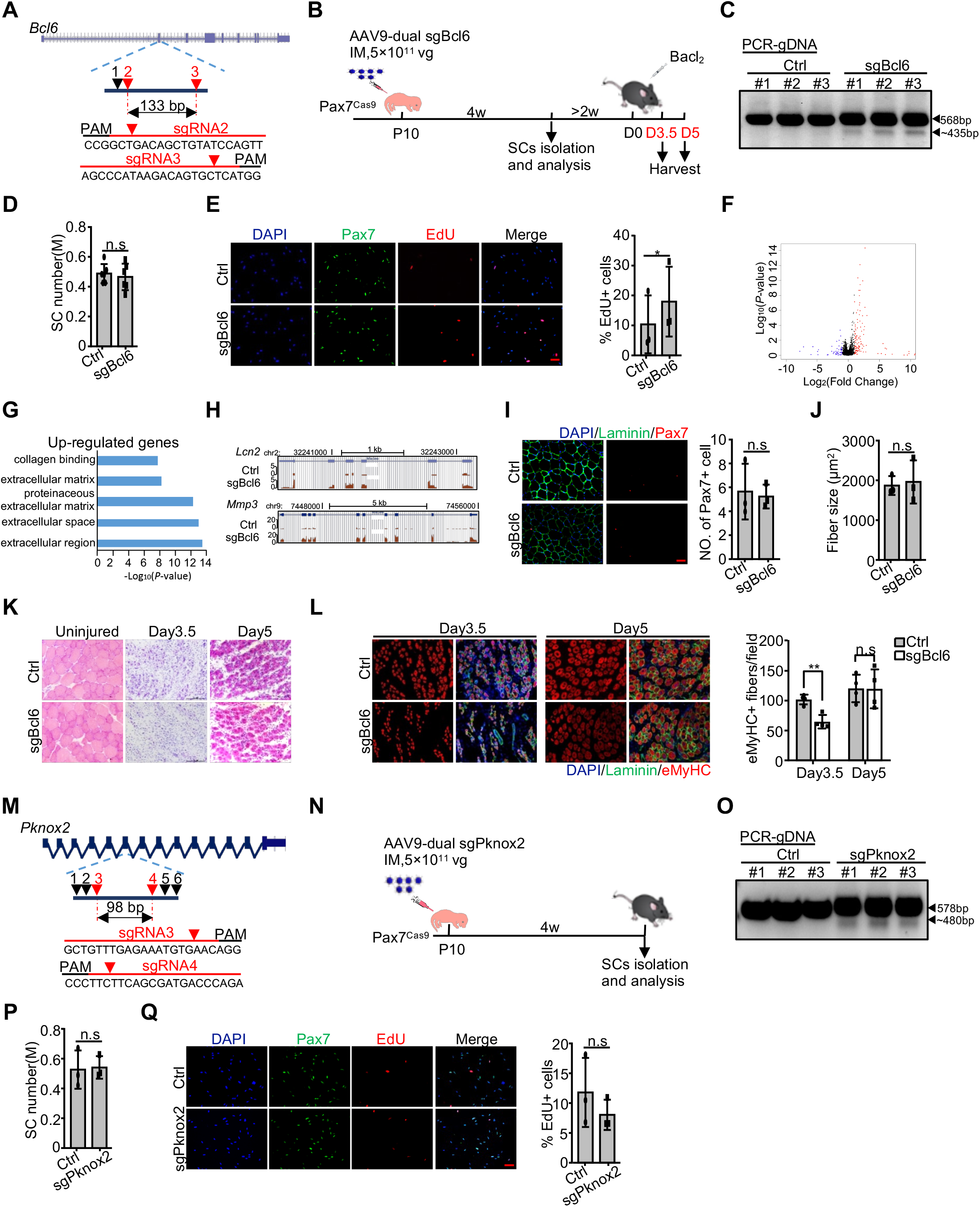
CRISPR/Cas9/AAV9 mediated genome editing of *Bcl6* and *Pknox2* led to distinct effects on SC activation and muscle regeneration. (**A**) Design and selection of sgRNAs targeting *Bcl6* locus. SgRNA2 and sgRNA3 were selected to delete a 133 bp of exon 3. (**B**) Schematic illustration of the experimental design for *in vivo* genome editing of *Bcl6* locus followed by analysis of its effect on SCs and muscle regeneration. Pax7_Cas9_ mice were administrated with high dose of control or AAV9-dual sgBcl6 virus at P10 and SCs were isolated four weeks later for analyses. To assay for muscle regeneration, BaCl_2_ was intramuscularly injected at the TA region of one leg at least six weeks after AAV9 virus administration. The injected muscle was harvested 3.5 or 5 days post injury and subject to analysis. (**C**) Genomic DNAs were isolated from the above sorted SCs and PCR analysis was performed to test the cleavage efficiency. Wide type (568 bp) and cleaved fragments (~435 bp) are shown by arrowheads. (**D**) The number of SCs isolated from the above control or dual sgBcl6 virus treated mice (n = 6 in each group). M: million. (**E**) The above FISCs were labeled with EdU for 24 hrs and the percentage of EdU positive cells was quantified (n = 3 in each group). Scale bar, 50 μm. (**F**) RNAs were isolated from the above SCs cultured for 24 hrs and subject to RNA-seq analysis. Differentially expressed genes were identified with blue and red dots representing down- or up-regulated genes, respectively. (**G**) GO analysis of the above up-regulated genes was performed and the top five enriched items are shown in y axis. X axis shows the -log10[*P* value] of the GO terms. (**H**) Genomic snapshots showing two examples of significantly up-regulated genes in sgBcl6 vs. ctrl treatment. (**I**) Immunostaining of Pax7 (red) and Laminin (green) were performed on the uninjured contralateral TA muscle (Left). The number of Pax7 positive cells per 100 fibers was counted (Right) (n = 3 in each group). Scale bar, 50 μm. (**J**) Average fiber size of the above TA muscle was quantified (n = 3 in each group). (**K**) H&E staining was performed 3.5 or 5 days post BaCl_2_ injection. Uninjured TA muscle was used as control (Left). Scale bar, 100 μm. (**L**) Immunostaining of eMyHC (red) and Laminin (green) on the above TA muscle sections (Left) and the number of eMyHC+ myofibers was counted (Right) (n = 4 in each group). Scale bar, 50 μm. (**M**) Design and selection of sgRNAs targeting *Pknox2* locus. SgRNA3 and sgRNA4 were selected to delete a 98 bp of exon 6. (**N**) Schematic illustration of the experimental design for *in vivo* genome editing of *Pknox2* locus and analysis of its effect on SCs. Pax7_Cas9_ mice were administrated with high dose of control or AAV9-dual sgPknox2 virus at P10 and SCs were isolated four weeks later for analysis. (**O**) Genomic DNAs were isolated from the above sorted SCs and PCR analysis was performed to test the cleavage efficiency. Wide type (578 bp) and cleaved fragments (~480 bp) are shown by arrowheads. (**P**) The number of SCs isolated from the above control or dual sgPknox2 virus treated mice (n = 3 in each group). M: million. (**Q**) The above FISCs were labeled with EdU for 24 hrs and the percentage of EdU positive cells was quantified (n = 3 in each group). Scale bar, 50 μm. All the bar graphs are presented as mean ± s.d. *P<0.05, **P<0.01. ns, no significance.

When examining the potential defects in muscle regeneration (Figure 7B), we found that AAV9-dual sgBcl6 injected mice displayed no overt changes of body weight or muscle weights (Supplementary Figure S6F). The number of Pax7+ cells was not affected neither (Figure 7I) and the average fiber size presented no abnormality (Figure 7J). These findings indicated that Bcl6 may not play a role in maintaining SC homeostasis. After BaCl_2_ induced injury, we failed to observe evident alteration in the degree of muscle regeneration in dual sgBcl6 vs. Ctrl group as assessed by H&E staining (Figure 7K and Supplementary Figure S6G), immunostaining of eMyHC expression (Figure 7L) and the measurement of fiber size 5 days post-injury (Supplementary Figure S6H). However, 3.5 days after injury, a reduced number of eMyHC+ cells was noticed which appeared to be contradictory to the repressive function of Bcl6 in SC activation (Figure 7L). We reasoned Bcl6 may play pleiotropic roles in SC lineage progression beyond promoting SC activation. Indeed, previous study has demonstrated that Bcl6 promotes myoblast differentiation by inhibiting apoptotic cell death(46). Altogether, the CRISPR/Cas9/AAV9 mediated genome editing has allowed us to dissect Bcl6 function in SCs *in vivo*.

Lastly, when applying the *in vivo* editing system on *Pknox2* using two sgRNAs targeting exon 6 (Figure 7M-N and Supplementary Figure S6I-J), evident excision of nearly 98 bp of *Pknox2* locus in SCs with high dose of AAV9-dual sgPknox2 virus (Figure 7O and Supplementary Figure S6K) did not appear to cause any alteration in SC homeostasis or activation (Figure 7P and Q). This is contrary to the finding from using siRNAs *ex vivo* (Figure 1G).

## DISCUSSION

In this study, we defined distinct lists of key TFs in SC lineage progression particularly in the early stage of SC quiescence and activation. Furthermore, with the help of the Cre-dependent Cas9 knockin mouse and AAV9 mediated sgRNA delivery, we created a muscle specific *in viv*o genome editing system. Using *MyoD* locus as a proof of concept, we demonstrated that this CRISPR/Cas9/AAV9 system can efficiently introduce mutagenesis at target TF locus and reduce its expression in SCs. Further application of the system on *Myc, Bcl6* and *Pknox2* revealed their distinct functions in the early stage of SC activation and damage induced muscle regeneration.

Signal pathways regulating SC quiescence and activation have been widely studied(1), our knowledge of key TFs involved in these stages however is very limited due to the lack of effective method to predict the stage-specific key TFs. Leveraging the recent findings showing that the expression of celltype-specific key TFs are generally regulated by SEs and bind to SE regions associated with their own and other key TFs(16,18,44), we were able to identify lists of key TFs in QSCs and ASCs respectively (Figure 1B), including the known SC fate decision factors like Pax7 and MyoD. Although we attempted to validate the predicted lists of key TFs using siRNA oligos, the *ex vivo* system prevents accurate analysis of SC quiescence and early activation stages. A prominent advantage of our CRISPR/Cas9/AAV9 system is that it allows for manipulation of SC genomes *in situ* without requiring cell isolation or culture, thereby making it possible to study the impact on SC quiescence, homeostasis in their native niche and early activation upon damage.

Despite recent demonstrations that the Cre-dependent Cas9-expressing mouse can be used to facilitate *in vivo* genome editing, ours represents the first effort of applying it on SCs or any somatic stem cells *in vivo*. On the paradigm of *MyoD* locus, it exhibited tremendous success in achieving nearly complete knockout of MyoD protein even at the lowest dose of AAV9 (0.2×10_11_ vg/mouse) carrying a single sgRNA. Consistent with prior reports(39,41,62), using dual sgRNAs resulted in an even higher editing at *MyoD* locus compared to single sgRNA. The attempt on *Myc* locus was also successful with Myc protein largely depleted. A high dose of AAV (5 × 10_11_ vg/mouse) was needed in this case, indicating the optimal AAV dosage is locus dependent. The editing efficiency on *Bcl6* or *Pknox2* was however not high despite the use of high dose of virus and dual sgRNAs, suggesting that the editing efficiency is locus dependent on the inherent quality of different protospacer sequences(61) or the chromatin state of each locus(68).

Phenotypes elicited by editing the three key TFs were highly diversified, revealing their previously uncharacterized functions in modulating early stages of SC activation. Loss of Myc led to evident decrease of SC activation thus delaying damage induced regenerating process. Considering its induction in FISCs vs. QSCs (Supplementary Figure S2A), Myc is probably one of the TFs key to exit of quiescence and entry of the first cell cycle, which warrants future endeavors in detailed characterization of its functional mechanism in SC and muscle regeneration. Loss of Bcl6 on the other hand slightly promoted SC activation. Although editing of *Pknox2* did not cause obvious change in SC activation, we cannot rule out the possibility that it may function in other stages of SC lineage progression.

Overall, the CRISPR/Cas9/AAV9 system manifests several advantages. First and the most prominent, compared with commonly employed transgenic and gene knockout-based models that necessitate significant investment of both time and resources, our approach can accelerate the pace at which gene function and interactions can be interrogated in SCs *in vivo.* Ultimately, this system can be adapted to enable rapid and direct *in vivo* screening of candidate and unknown gene targets suspected to specifically influence SC phenotypes. The ability of achieving multiplex genetic perturbations using the Cas9 mouse also enables the possibility of interrogating multigenic effects. As multigene interactions play a critical role in SC activities like in virtually any biological process, in the future, it will be of our main interest to further develop this system to allow for simultaneous study of small sets of genes or even high-throughput *in vivo* genetic screening, for example, by combining with barcode labelling and single cell RNA-seq, this system can be employed to simultaneous study multiply genes modulating SC function and uncover novel biology. Similar approach has been used to identify functional suppressors in glioblastoma by combining CRISPR/Cas9/AAV mediated *in vivo* genome editing with targeted capture sequencing(69). In addition to coding genes, it is also possible to interrogate non-coding genome such as enhancers and lncRNAs, therefore further expanding the application of this system. For example, in a recent study(70), we have applied it to edit an eRNA in skeletal muscle which facilitated the investigation of the eRNA function in regulating myoblast differentiation.

Of note, another interesting fact is that Cas9 expression is present at birth in Pax7_Cas9_ mouse and the administration of AAV9 virus at P10 enables the investigation of postnatal myogenesis. Normally, juvenile SCs emerge about two days before birth in mice and continue to proliferate until about 12 days after birth to form adult muscle through undergoing postnatal myogenesis(71). Quiescent SCs appear at about two weeks after birth to build the SC pool(59). Genome editing at P10 will allow us to study the impact on SC pool establishment and homeostatic maintenance until adult period. For example, editing *MyoD* locus led to evident elevation of the number of FISCs isolated four weeks later, suggesting that MyoD loss in postnatal stage may have affected SC establishment. Editing *Myc, Bcl6* or *Pknox2,* nevertheless, did not cause such alteration.

One interestingly caveat is AAV9 administration may also edit muscle fibers since Cas9 is highly expressed in fibers as well. We attempted to solve this problem by using inducible Pax7_ER-Cas9_ mice that will allow SC restricted Cas9 expression but the results were not promising at the stage. Second, it should be kept in mind that this system does not generate a uniform gene knockout mouse. Although the transduction efficiency appeared to be close to 100%, the editing efficiency per SC and the editing events largely varied, leading to mosaic mutagenesis. Mosaic editing for some genes, for example, *Pknox2*, may not be sufficient to elicit any phenotypical alterations, in which case traditional knockout models may be needed to confirm the findings.

Lastly, it is necessary to stress that our attempts of editing quiescent SCs was unsuccessful. It is believed that QSCs cannot be edited *in vivo* owning to the limited transduction efficiency of AAV(25). Our strategy of administration of AAV at P10 allowed for efficient transduction when SCs are acceptable for AAV9 transduction. Still, no editing was achieved after Tmx induction of Cas9 expression. It has been reported that the repair of radiation-induced DSBs in non-proliferating SCs is not only efficient but also accurate depending on the key NHEJ factor DNA-PKcs to maintain genome integrity through direct ligation of the DSBs(72). It is thus possible that Cas9 may have cut the sites in QSCs, but the caused double-strand breaks were ligated precisely without any mutations at target sites. If so, the AAV9-dual sgRNA system should induce mutagenesis by deleting a DNA fragment, which however was not successful in our attempts on *MyoD, Myc,* and *Bcl6* loci. We thus suspect that the failure of the editing in QSCs may be caused by the condensed chromatins(73) that inhibited the function of Cas9 as recent studies have demonstrated that Cas9 prefers to exert its function at open chromatin regions(68,74).

## Author contributions

Conceived and designed the experiments: H.W., H.S., L.H. and Y.D. Performed the experiments: L.H., Y.Z., K.K.S., and Y.L. Analyzed the data: Y.D., X.L.P., and J.Y. Wrote the paper: H.W., L.H. and Y.D. Reviewed and edited the manuscript: H.W., H.S., L.H. and Y.D.

## FUNDING

Research Grants Council (RGC) of the Hong Kong Special Administrative Region (Project Code: 14100415, 14133016, 14106117, and 14100018 to H.W.; 14102315 and 14116918 to H.S.); CUHK Focused Innovations Scheme: Scheme B to H.S. (Project Code: 1907307); National Natural Science Foundation (NSFC) of China (Grant No: 31871304 to H.W.); NSFC/RGC Joint Research Scheme from RGC (Project Code: N_CUHK413/18 to H.S.). Direct Grant from CUHK (Project Number: 2017.049 to H.W. and 2017.009 to H.S.).

## Conflict of interest statement

The authors declare that they have no conflict of interest.

## Notes

### Competing Interest Statement

The authors have declared no competing interest.

### Summary of Updates

The order of the authorship has been changed.

